# IL-4 downregulates BCL6 to promote memory B cell selection in germinal centers

**DOI:** 10.1101/2023.01.26.525749

**Authors:** Laila Shehata, Christopher D. Thouvenel, Brian D. Hondowicz, Lucia A. Pew, David J. Rawlings, Jinyong Choi, Marion Pepper

## Abstract

Germinal center (GC)-derived memory B cells (MBCs) are critical for humoral immunity as they differentiate into protective antibody-secreting cells during re-infection. GC formation and cellular interactions within the GC have been studied in detail, yet the exact signals that allow for the selection and exit of MBCs are not understood. Here, we show that IL-4 signaling in GC B cells directly downregulates BCL6 via negative autoregulation to release cells from the GC program and promote MBC formation. This selection event requires additional survival cues and can therefore result in either GC exit or death. We demonstrate that both increasing IL-4 bioavailability or limiting IL-4 signaling disrupt MBC selection stringency. In this way, IL-4 control of BCL6 expression serves as a tunable switch within the GC to tightly regulate MBC selection and affinity maturation.

## Introduction

Germinal centers (GCs) are highly organized structures found within B cell follicles. In the GC, activated B cells proliferate, diversify their B cell receptors, and are selected to mature into memory cells that provide humoral protection from disease^1^. Temporally and spatially distinct and complex interactions between B cells and other cells in the GC including CD4^+^ T follicular helper (Tfh) cells and follicular dendritic cells (FDCs) influence the initiation, maintenance, and dissolution of GCs. While many of these interactions have been well-characterized, many others, including how affinity-matured memory B cells (MBCs) are selected to exit the GC, remain poorly defined^2–9^. Specifically, the mechanisms underlying the CD4^+^ T cell-mediated selection process that leads to the downregulation of GC B cell-defining transcriptional programs and concomitant upregulation of transcriptional programs associated with long-lived memory B cell formation are unknown^10–12^. It is well accepted that BCL6, a repressive transcription factor, actively maintains the GC and must be downregulated for MBCs to form, but the signals that initiate the downregulation of BCL6 have not been identified^13,14^. Understanding how this process takes place is critical for generating high affinity memory B cell formation for optimal vaccine development or for targeting this process therapeutically during autoimmunity.

Interleukin-4 (IL-4) is a key T cell-derived molecule that has been attributed with conflicting roles in the regulation of the GC^15,16^. Early after the initiation of an anti-viral response, IL-4 produced by natural killer T cells is able to stimulate GC formation by promoting B cell expression of BCL6^15,17,18^. BCL6 acts via repressor-of-repressor circuits to modulate transcriptional programs that keep B and Tfh cells tethered in the GC and prevent them from terminally differentiating^14,19,20^. BCL6 also suppresses the DNA damage sensor ATR, thereby acting as an anti-apoptotic factor to keep GC B cells alive as they undergo rounds of DNA damage-prone somatic hypermutation^21,22^. Yet how subsequent IL-4 production by CD4^+^ Tfh and GC Tfh cells impacts BCL6 expression, GC maintenance, and memory formation is unclear. Recent work from Duan and colleagues demonstrated that excess availability of IL-4 could reduce memory B cell formation and reduce selection stringency, although the underlying mechanisms were not defined^16^. Herein we have sought to define how IL-4 functions in the GC and to determine if it could influence the selection or exit of MBCs through its regulation of BCL6. We demonstrate that while IL-4 promotes BCL6 in naive and activated B cells to encourage GC formation, it can also induce the negative autoregulation of BCL6 in a GC B cell-intrinsic manner. This IL-4-mediated downregulation of BCL6 initiates a selection event, allowing pre-memory GC B cells that can obtain additional survival signals to exit the GC, whereas those that cannot undergo cell death.

## Results

### IL-4 can be produced by GC Tfh cells and perceived by GC B cells during Plasmodium infection

To better understand how CD4^+^ T cell-derived IL-4 can impact B cell differentiation during infection, we first characterized the identity and kinetics of IL-4-producing CD4^+^ T cells specific for *Plasmodium*-derived antigens during a GC response. To accomplish this, KN2 (knock-in huCD2) mice that express cell surface-associated huCD2 protein under control of the IL-4 promoter were infected with 10^6^ *Plasmodium yoelii*-GP66 (*P.y*-GP66)-infected red blood cells (iRBCs). We examined IL-4 production from days 6-18 after *Plasmodium* infection as our previous work demonstrated that this is a critical window for *Plasmodium*-specific B cell differentiation, encompassing the initiation and propagation of the GC as well as early memory formation^23^. IL-4 producing GP66-specific CD4^+^ T cells were identified using previously published tetramer enrichment and flow cytometric techniques^24^. In response to *Plasmodium*, the number of IL-4-producing T cells increased throughout the acute stage of infection, as previously shown in influenza and helminth infections^18,25,26^ (**Figure S1A and S1B**). Of interest, although Tfh cells contributed to early IL-4 production, the majority of IL-4-producing CD4^+^ T cells identified were GC Tfh (**Figure S1B**). This indicates that in a Type 1 immune response to infection, the bulk of CD4^+^ T cell-derived IL-4 is produced by GC Tfh cells after the GC has formed, suggesting contributions beyond solely early GC formation.

We generated B cell tetramers containing the *Plasmodium* blood-stage antigen Merozoite Surface Protein-1 (MSP1) to characterize IL-4 receptor alpha chain (IL4Rα; CD124) expression on responding B cells over the course of infection^23^. There was a large population of IL4Rα^+^ MSP1-specific CD38^+^GL7^-^ B cells in uninfected mice (day 0), suggesting naive B cells can receive IL-4 signals early in infection (**Figure S1C and S1D**). IL4Rα expression remained high on MSP1-specific CD38^+^GL7^-^ as well as GL7^+^ B cells throughout acute *P.y*-GP66 infection (**Figure S1D**). Overall, this demonstrates that B cells are capable of perceiving Tfh- and GC Tfh-derived IL-4 both prior to and within an ongoing GC.

### IL-4 has opposing effects on the B cell response over the course of infection

The kinetics of CD4^+^ GC Tfh production of IL-4 in conjunction with GC B cell expression of IL4Rα suggested a role for IL-4 in the GC that perhaps went beyond solely GC entry. We therefore asked how *in vivo* administration of exogenous IL-4 at various time points affects B cell development after *Plasmodium* infection. In an effort to maximize IL-4 bioavailability, we utilized complexed IL-4 (IL4C; recombinant IL-4 complexed to anti-IL-4 antibody) to prolong the half-life of the cytokine *in vivo*^27^, a strategy commonly used to assess *in vivo* exogenous IL-4 signaling^16,18^. We infected C57BL/6 (WT) mice with 10^6^ *Plasmodium chabaudi* (*P.ch*)-iRBCs and administered IL4C or phosphate buffered saline (PBS) intravenously on the first 3 days of infection to determine if early IL-4 enhances CD38^-^GL7^+^ GC B cell differentiation as seen in influenza and other infections^18,27^ (**Figure 1A**). We observed a significant increase in the number and frequency of MSP1-specific GC B cells by day 8 post-infection in mice treated with IL4C compared to untreated controls, suggesting that this role for early IL-4 in promoting GCs is common to multiple infections^18^ (**Figure 1B and 1C**).

**Figure 1.**
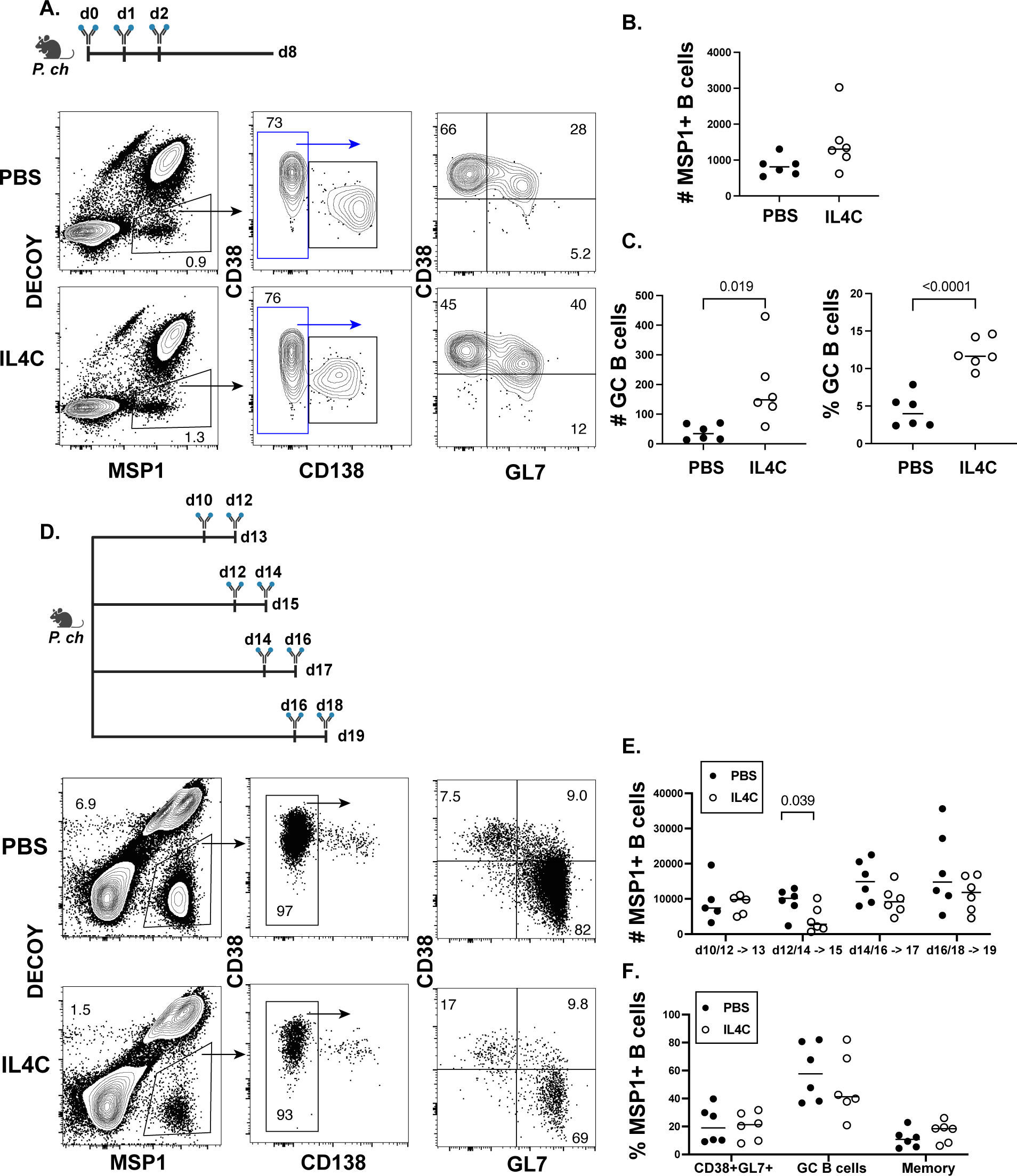
The effect of IL-4 on the antigen-specific B cell response varies over the course of infection. (**A**) WT mice were infected with *P.ch* and treated with PBS or IL4C on days 0, 1, and 2 post-infection. Representative flow cytometry plots of MSP1-specific cells at 8 days post-infection from mice treated with PBS (top) or IL4C (bottom), with further gating on non-plasma cells (CD138^-^) and GC B cells (CD38^-^GL7^+^) as defined in Figure S1. (**B**) Total MSP1-specific B cells in each treatment group at day 8. (**C**) Number (left) and frequency (right) of MSP1-specific GC B cells in each treatment group at day 8. Data are combined from two independent experiments with 6 mice per group. (**D**) WT mice were infected with *P.ch* and treated with IL4C or PBS over two-day windows and MSP1-specific B cell responses were analyzed the following day, as indicated. Representative flow cytometry plots from mice treated with PBS (top) or IL4C (bottom) on days 12 and 14 and analyzed on day 15. (**E**) Number of MSP1-specific B cells at each time point analyzed. (**F**) Frequency of CD38^+^GL7^+^, GC B cells, and MBCs (CD38^+^GL7^-^ CD73^and/or^CD80^+^) in each treatment group at day 15, following PBS or IL4C treatment on days 12 and 14. Data are combined from two independent experiments with 5-6 mice per group. See also Figures S1 and S2.

We next asked if IL4C administration to *P.ch*-infected WT mice during an ongoing GC had a similar effect. Mice were administered PBS or IL4C in staggered three-day windows following GC formation (day 10) and the MSP1-specific B cell response was analyzed one day following the last treatment (**Figure 1D**). Although there was no significant impact on the B cell response when mice received IL4C on days 10 and 12, there was a surprising loss of MSP1-specific B cells when IL4C was administered on days 12 and 14 (**Figure 1E**). A similar trend was observed for the later IL4C treatment windows as well. Importantly, MSP1-specific cells in IL4C-treated mice did not have higher plasmablast numbers, BLIMP1 expression, or anti-MSP1 serum IgG, indicating that IL-4 is not diverting MSP1-specific B cells to a plasmablast fate (**Figure S2A-S2C**). Thus, although early IL-4 encourages GC formation and the ensuing antigen-specific GC B cell expansion, IL-4 restrains the B cell response once GCs have developed. Of interest, in mice treated with IL4C on days 12 and 14 that experienced the greatest loss of MSP1-specific B cells, there were no significant changes in the frequencies of any one specific B cell subset examined (**Figure 1F**), suggesting that IL-4 may be impacting and restricting the GC process, which could affect multiple subsets equally.

### IL-4 downregulates BCL6 and enhances expression of CD80

To test if IL-4 was restricting the GC, we next imaged spleens to visualize GCs in mice treated with IL4C on days 12 and 14. Interestingly, we found that GCs in IL4C-treated mice were smaller in size, supporting our hypothesis that IL-4 can constrain an ongoing GC response (**Figure S3A and S3B**). This could reflect an effect of IL-4 on GC precursors, GC B cells, exiting GC B cells, or all three B cell populations. Cells transitioning through a CD38^+^GL7^+^ state (see model in **Figure 2A**) include both GC precursors and pre-memory, exiting GC B cells as shown in analyses of B cell populations in UBP-2A-Fucci mice (which report distinct phases of cell cycle) as well as B cells responding to infection with lymphocytic choriomeningitis virus (Armstrong strain; LCMV_arm_)^28–30^. However, the markers used to identify these pre-memory cells thus far are proteins that are either also expressed on GC precursors (CCR7, CD62L) or are also expressed prior to and within the GC (Ephrin B1)^29,30^. This makes it difficult to accurately define B cells exiting the GC by flow cytometry. We therefore sought to identify a marker of GC-exiting pre-memory B cells that is not expressed on GC precursors or GC B cells but is upregulated on MBCs. We hypothesized that within the CD38^+^GL7^+^ “GC transit” B cell population, differing expression of BCL6 and the co-stimulatory molecule CD80 could be used to distinguish pre-GC B cells from post-GC B cells. Activated B cells upregulate BCL6 very early after immunization, and GC transit B cells show increasing BCL6 expression as the GC develops^31^ (**Figure 2B and 2C**). In order to form memory B cells, GC B cells must downregulate BCL6^14,32,33^. This releases the repression of numerous migratory and functional programs, including allowing the expression of CD80^34^. Indeed, we did not observe CD80^+^ cells in the GC when BCL6 is most highly expressed, yet a distinct population of CD80^+^BCL6^-^ GC transit B cells emerges after cells begin to downregulate BCL6 and form MBCs 10 days after *Plasmodium* infection^35,36^ (**Figure 2B and 2C**). We can therefore identify “entering” GC transit cells as BCL6^+^CD80^-^CD38^+^GL7^+^ B cells and “exiting” GC transit cells as BCL6^-^CD80^+^CD38^+^GL7^+^ GC B cells.

**Figure 2.**
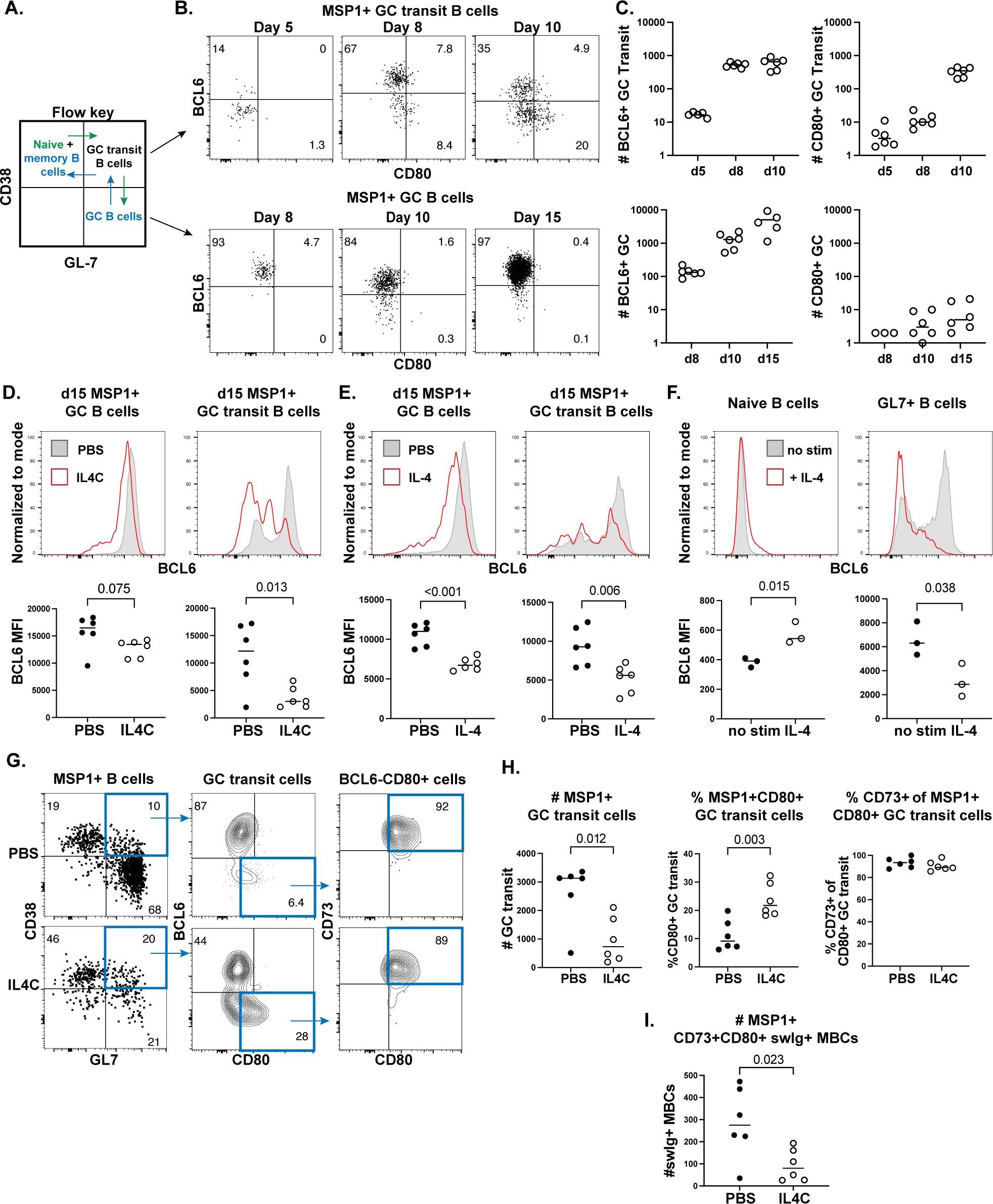
IL-4 downregulates BCL6 during an ongoing GC. (**A**) Flow key indicating the identification of naive and memory B cells, GC transit B cells, and GC B cells by CD38 and GL7 expression. (**B**) Representative flow plots and (**C**) quantification of BCL6 and CD80 expression in MSP1^+^ GC transit B cells (bottom) and GC B cells (top) from *P.ch*-infected WT mice on the indicated days post-infection. Data are combined from two independent experiments with 6 mice per group. Representative histograms (top) and quantified mean fluorescent intensity (MFI) of BCL6 expression (bottom) in GC B cells and GC transit B cells at day 15 in mice treated with (**D**) PBS (grey) or IL4C (red) on days 12 and 14 post-*P.ch* infection or (**E**) PBS or non-complexed IL-4 on days 12, 13, and 14 post-*P.ch* infection. Data in (D) and (E) are each combined from two independent experiments with 6 mice per group. (**F**) Naive or GL7^+^ B cells were isolated from naive or *P.ch*-infected murine spleens, respectively, and cultured with agonistic anti-CD40 antibody with or without non-complexed IL-4. Representative histograms and quantified MFI of BCL6 expression after 24h. Data are combined from three independent experiments, with each symbol representing the average of three technical replicates from each experiment. (**G**) Representative flow plots and (**H**) quantification of GC transit B cell expression of CD80 and CD73 in mice treated with PBS or IL4C. (**I**) Number of MSP1^+^swIg^+^CD73^+^CD80^+^ MBCs. Data are combined from two independent experiments with 6 mice per group. See also Figure S3.

To test if and how IL-4 could be impacting these populations, we analyzed BCL6 and CD80 expression on MSP1^+^ B cells by flow cytometry after IL4C treatment. In opposition to the current paradigm that IL-4 promotes BCL6, we observed an insignificant decrease in BCL6 expression in GC B cells as well as a significant loss of BCL6 in GC transit B cells in mice treated with IL4C on days 12 and 14 compared to controls (**Figure 2D**). To confirm that this was not unique to an IL4C signal as opposed to non-complexed IL-4, we also treated mice with non-complexed IL-4 on days 12-14. Both the loss of MSP1-specific cells noted in Figure 1 as well as the loss of BCL6 expression were recapitulated with IL-4 alone, demonstrating that this phenotype is IL-4-mediated and not dependent upon signaling alterations due to complexing IL-4 with antibody (**Figure 2E; Figure S3C**).

Our data suggested that IL-4 signaling may differentially impact B cells at different stages of differentiation during an immune response. To test this hypothesis directly, we isolated naive splenic B cells from naive mice or GL7^+^ splenic B cells from mice at least 12 days post-*Plasmodium* infection and cultured them with an agonistic anti-CD40 antibody (1 ug/mL) to promote *in vitro* GC B cell survival^37,38^. Both cell types were cultured in the presence or absence of non-complexed recombinant IL-4 (10 ng/mL) for 24 hours. Naive B cells analyzed after 24h culture with anti-CD40 and IL-4 had significantly higher BCL6 expression than those cultured with anti-CD40 alone, consistent with previous literature^17^ (**Figure 2F**). However, GL7^+^ B cells cultured in the presence of anti-CD40 and IL-4 lost BCL6 expression (**Figure 2F**), mirroring the *in vivo* phenotype observed in mice treated with IL4C or IL-4 **(Figure 2D and 2E)**. This indicates that IL-4 can have opposing effects on BCL6 expression in B cells at various stages of differentiation, perhaps contingent on the level of BCL6 already present in the cell.

While the reduction in BCL6^+^ GC transit B cells could be indicative of a loss of GC precursors, it corresponded to an increase in the frequency of CD80^+^ GC transit B cells, suggesting that IL-4 may be downregulating BCL6 in GC B cells, resulting in increased GC exit (**Figure 2G and 2H**). We further analyzed expression of the ectonucleotidase CD73, used to identify functional memory B cells that have received CD4^+^ T cell help^23,35,36,39^. Nearly all of the CD80^+^ GC transit B cells in both PBS- and IL4C-treated mice co-expressed CD73, further supporting the idea that these are indeed exiting pre-memory B cells (**Figure 2G and 2H**). Surprisingly however, despite the proportional increase in exiting CD73^+^CD80^+^ GC transit B cells, IL4C-treated mice exhibited a specific loss of GC-derived class-switched (IgM^-^IgD^-^; swIg^+^) CD73^+^CD80^+^CD38^+^GL7^-^ MBCs (**Figure 2I; Figure S3D**). Thus, although IL-4 appears to be increasing GC exit into the GC transit pool, the overall loss of MSP1-specific B cells and of GC-derived MBCs suggests that the IL-4-mediated downregulation of BCL6 may not be the only signal required for long-lived MBC formation.

### IL-4 directly downregulates BCL6 on GC B cells via negative autoregulation

We sought to understand both how IL-4 was regulating BCL6 expression in GC B cells and why this may not be enough to generate long-lived MBCs. Although our primary B cell culture data suggested that IL-4 is acting directly on B cells to downregulate BCL6, we aimed to confirm this *in vivo* using mice in which IL4Rα is conditionally deleted from GC B cells. S1PR2 is upregulated as B cells enter the GC and acts as a tethering protein to maintain B cell positioning within the GC^40^. S1PR2creERT2^+/−^TdTomato^flox^ mice have been shown to accurately label GC B cells and fate map GC-derived MBCs^5^. We therefore crossed S1PR2creERT2^+/−^ Tdtomato^flox^ mice to IL4Rα^flox^ mice to allow for the conditional deletion of IL4Rα from GC B cells after the GC has formed. We infected S1PR2creERT2^+/−^Tdtomato^flox^ IL4Rα^flox^ and their S1PR2creERT2^-/-^ littermates with *P.ch* and fed them tamoxifen chow beginning on day ten^5^. We then analyzed the MSP1-specific B cell response on day 20 to determine if IL4Rα-expressing GC-experienced cells were more likely to have downregulated BCL6 and exited the GC compared to IL4Rα-deficient cells, which cannot respond to IL-4 (**Figure 3A**). Using TdTomato as a fate reporter for S1PR2 expression, we observed that while >90% of MSP1-specific B cells were TdTomato^+^, only ∼50-70% of the cells within each B cell subset had successfully deleted IL4Rα (as seen by flow cytometric analysis of anti-IL4Rα staining), as has been previously described (**Figure 3A**)^16^. Interestingly, within the TdTomato^+^ cells from the S1PR2creERT2^+/−^ mice, the frequency of BCL6^+^ GC transit B cells was significantly higher in the IL4Rα^-^ population compared to the IL4Rα^+^ population (**Figure 3B**). This loss of BCL6 in IL4Rα^+^ GC transit B cells additionally corresponded to an increase in CD80^+^ GC transit B cells, further supporting that these CD80^+^ cells GC transit cells are GC-experienced, as they were fate-mapped with S1PR2 (**Figure 3B**). Together, these findings demonstrate that IL-4 is directly acting on GC B cells to downregulate BCL6.

**Figure 3.**
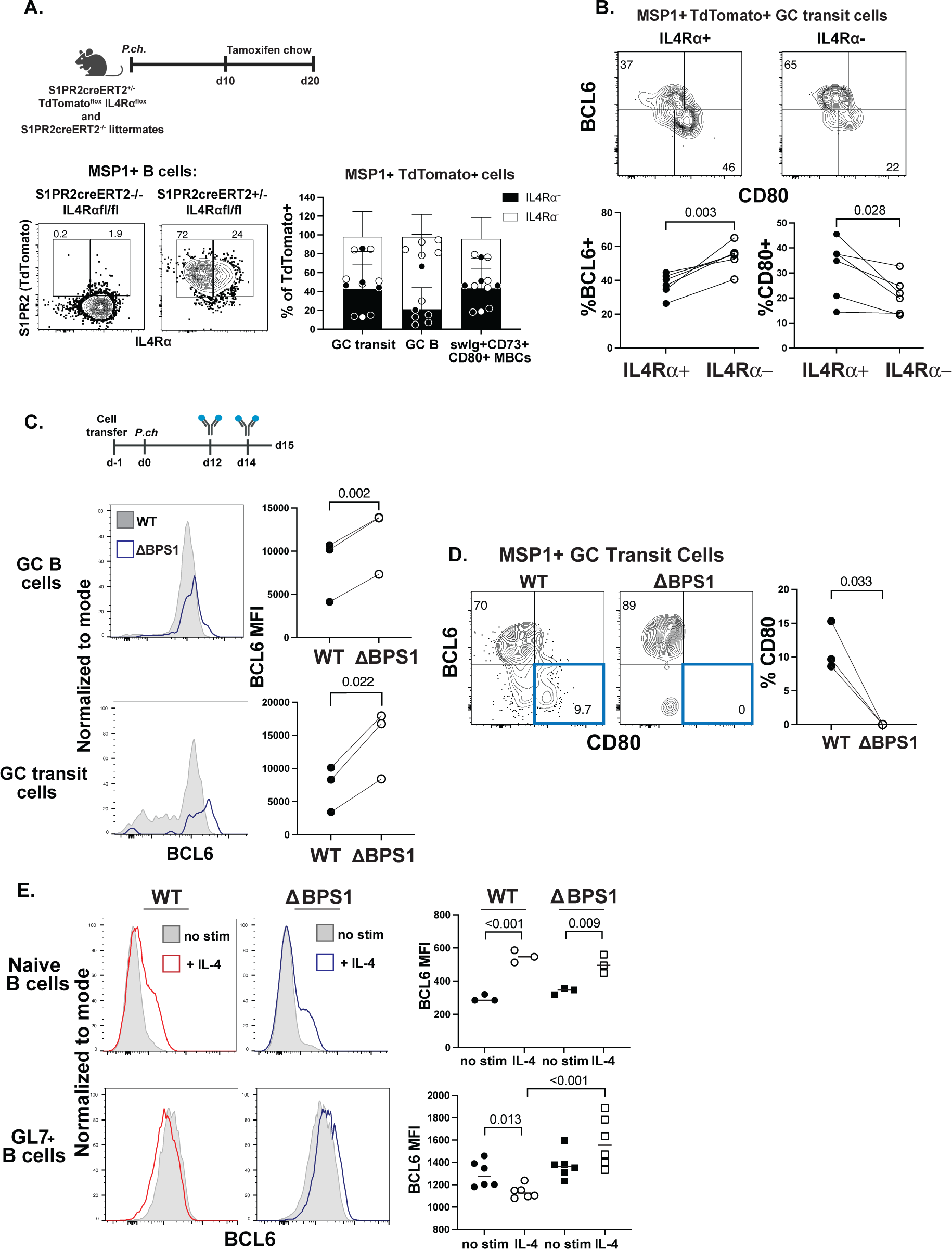
IL-4 signaling in GC B cells triggers BCL6 negative autoregulation. (**A**) S1PR2creERT2^+/−^TdTomato^flox^IL4Rα^flox^ mice and S1PR2creERT2^-/-^ littermates were infected with *P.ch* and fed tamoxifen chow beginning 10 days post-infection and analyzed on day 20. Representative flow plots and quantification of TdTomato^+^ cells that retained or successfully deleted IL4Rα shown on the bottom. IL4Ra gating based on FMO control, as shown in Figure S1. (**B**) Representative flow plots (top) and quantification (bottom) of BCL6 and CD80 expression within IL4Rα^+^ or IL4Rα^-^ MSP1^+^TdTomato^+^ GC transit B cells from S1PR2creERT2^+/−^ TdTomato^flox^ IL4Rα^flox^ mice. Data are combined from three independent experiments with 6 mice per group. (**C**) Splenic B cells were isolated from ΔBPS1 mice and transferred into WT hosts. The mice were infected with *P.ch*, treated with IL4C on days 12 and 14, and the MSP1-specific B cell response was analyzed on day 15. Representative histograms and quantification of BCL6 expression in WT (grey) and ΔBPS1 (blue) GC B cells and GC transit B cells on day 15. (**D**) Representative flow plots depicting BCL6 by CD80 expression in MSP1^+^ GC transit B cells and quantified frequency of CD80^+^ GC transit B cells. Data are from two independent experiments with three mice. (**E**) BCL6 expression on naive or GL7^+^ B cells isolated from naive or LCMV-infected murine spleens, respectively, and cultured with agonistic anti-CD40 antibody with or without non-complexed IL-4 for 24h. Data are from one experiment, with each symbol representing replicates from one (naive) or two pooled (GL7^+^) mice. See also Figure S4.

As it was now evident that expression of IL4Rα on GC B cells was required for the IL-4-mediated loss of BCL6 expression in GL7^+^ B cells, we next sought to understand how IL-4 was differentially affecting BCL6 in B cells. Important work in the oncology field and more recently in CD4^+^ T cells has demonstrated that BCL6 is able to negatively autoregulate its own expression, a process that is often disrupted in B cell lymphomas^19,41,42^. It was formally possible that while early IL-4 enhances BCL6 expression and GC entry early in an immune response, administration of IL-4 to GC B cells that express a high enough threshold of BCL6 may promote BCL6 to bind its own promoter and negatively autoregulate its own expression. We therefore sought to elucidate the specific signaling components that may be involved in this regulatory cascade. IL-4 can signal through both STAT6 and IRS-2, although its ability to promote BCL6 expression has been shown to depend upon the STAT6 binding site in the BCL6 promoter region^17,43,44^. To demonstrate a role for STAT6 signaling in the IL-4-mediated repression of BCL6, we treated *P.ch*-infected WT and STAT6-deficient (STAT6KO) mice with IL4C and analyzed BCL6 and CD80 expression in MSP1-specific GC transit B cells as above. Of interest, both GC B cells and GC transit B cells in both WT and STAT6KO mice were able to express equivalent amounts of BCL6, consistent with prior literature demonstrating that IL-21 and other molecules contribute to B cell expression of BCL6^15,45,46^ (**Figure S4A**). However, while IL4C treatment of WT cells led to BCL6 downregulation and CD80 upregulation as shown above, GC transit B cells that lack STAT6 did not alter their expression of BCL6 or CD80 in response to IL4C, demonstrating that this IL-4 driven process is STAT6-dependent (**Figure S4A and S4B**).

We next directly tested if the IL-4-STAT6 pathway was promoting BCL6 to engage its negative autoregulation by using ΔBPS1 mice, which have an 8-nucleotide deletion in the BCL6 promoter binding site 1 (BPS1) that prohibits BCL6 from binding to its own promoter^19^. We adoptively transferred splenic B cells isolated from CD45.2^+^ ΔBPS1 mice into CD45.1^+^ WT hosts, infected the hosts with *P.ch* and treated the mice with IL4C on days 12 and 14, prior to analysis of the MSP1-specific B cell response on day 15 (**Figure 3C; Figure S4C**). In both GC B cells and GC transit B cells, BCL6 expression remained significantly higher in the ΔBPS1 cells than in WT cells (**Figure 3C**). Additionally, the IL-4-mediated downregulation of BCL6 in WT GC transit B cells corresponded to an increase in CD80 expression, whereas CD80 was not upregulated in ΔBPS1 GC transit B cells after IL4C treatment (**Figure 3D**). We additionally used the same primary murine B cell culture setup as above to determine how non-complexed IL-4 affects both ΔBPS1 naive and GC B cells. We isolated naive or GL7^+^ B cells from WT or ΔBPS1 naive mice or mice 10 days post-LCMV_arm_ infection, respectively, and cultured them with an agonistic anti-CD40 antibody (1 ug/mL) in the presence or absence of IL-4 (10 ng/mL) for 24h. IL-4 similarly increased BCL6 expression in WT and ΔBPS1 naive B cells (**Figure 3E**). However, while WT GL7^+^ B cells downregulated BCL6 in response to IL-4, consistent with our findings in Figure 2, ΔBPS1 GL7^+^ B cells showed no significant change in BCL6 expression following culture with IL-4 (**Figure 3E**). In fact, BCL6 expression was significantly higher in ΔBPS1 GL7^+^ B cells cultured with IL-4 compared to WT GL7^+^ B cells. Together, these data show that IL-4 signaling via STAT6 leads to BCL6 negative autoregulation in GL7^+^BCL6^+^ B cells, enhancing GC exit via a CD38^+^GL7^+^CD80^+^ transition stage, yet these cells do not survive to become MBCs.

### IL-4 mediated cell death limits the formation of GC-derived MBCs

To better understand why the IL-4-mediated downregulation of BCL6 was not associated with enhanced MBC formation and survival, we began to dissect where along the path of MBC formation these cells were lost. We first interrogated any changes in GC cycling, as a disruption of positive selection of light zone (LZ) GC B cells into the dark zone (DZ) could explain the lack of swIg^+^ MBCs (Figure 2I). However, there were no changes in the proportion of GC B cells in the LZ or DZ, as measured by CXCR4 and CD86 expression, in response to IL4C treatment (**Figure S5A**). We confirmed this by examining the expression of c-myc, a transcription factor which is transiently upregulated as GC B cells migrate into the DZ^11,47^. We did not observe any changes in c-myc expression in GC B cells, BCL6^-^CD80^+^ GC transit B cells (exiting GC), or CD73^+^CD80^+^ memory B cells in WT mice infected with *P.ch* and treated with IL4C or PBS (**Figure S5B**).

To test if IL4C treatment was impacting the formation of memory precursors in the GC or their successful exit, we additionally analyzed expression of CD23 and BCL2 on the three populations described above. CD23 is upregulated on GC B cells that have recently received IL-4 and/or CD40L signals from T cells, and CD23^+^ GC B cells have been identified as memory B cell precursors^16,47^. IL4C treatment led to a significant increase in CD23^+^BCL6^-^CD80^+^ GC transit B cells, whereas GC B cells and MBCs had modest, but statistically insignificant increases in CD23 expression in response to IL4C (**Figure 4A**). This suggests that IL-4 is promoting memory B cell differentiation, and is consistent with our hypothesis that the CD80^+^ GC transit B cells have recently left the GC in response to IL-4 signaling. We also stained for BCL2, an anti-apoptotic protein that is critical for MBC survival^10,33,48,49^. BCL2 is highly expressed by naive B cells, is downregulated upon GC entry, and is upregulated again as GC B cells form MBCs^48^. Importantly, it can repress BCL6, and it has been suggested that this antagonism is mutual^4,33^. Interestingly, GC B cells and MBCs from mice treated with IL4C had a striking loss of BCL2 compared to control-treated mice, while BCL2 expression in BCL6^-^CD80^+^ GC transit B cells was unchanged between groups (**Figure 4B**). However, ∼80% of MBCs were still BCL2^+^, suggesting the vast majority of these cells are still viable (**Figure 4B**). Together, these data suggest that IL-4 downregulates BCL6 in GC B cells to promote the exit and formation of MBCs, but is insufficient to upregulate BCL2, which could replace BCL6 as an anti-apoptotic signal. This may result in an increase in cells dying while exiting the GC due to a lack of BCL2 upregulation within the GC, which would explain the numerical loss of GC transit B cells and GC-derived MBCs following IL4C treatment (Figure 2H and 2I).

**Figure 4.**
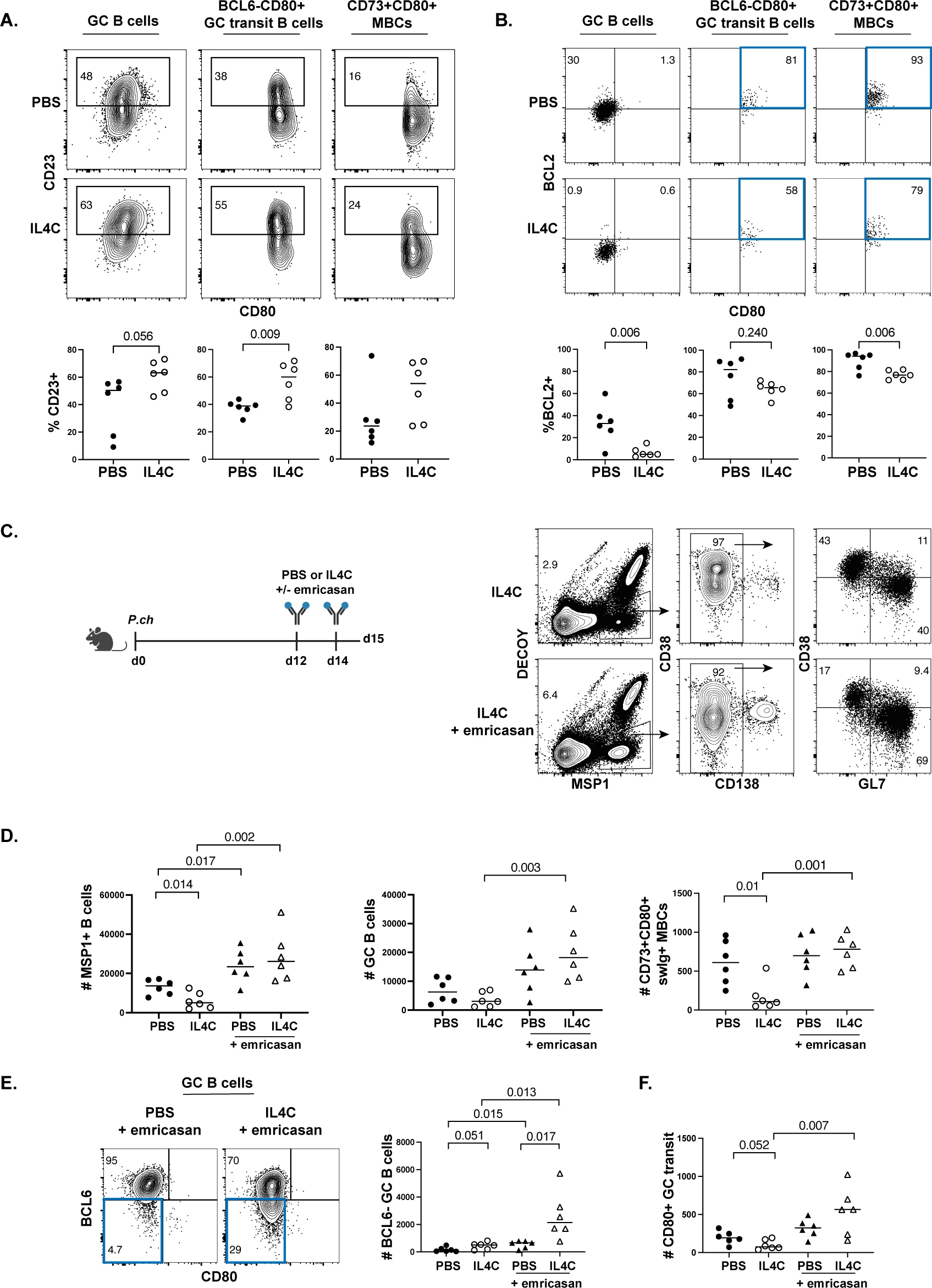
Excess IL-4 leads to increased GC B cell death and loss of MBCs. WT mice were infected with *P.ch* and treated with IL4C or PBS on days 12 and 14, and the MSP1-specific B cell response was analyzed on day 15. Representative flow plots and quantification of MSP1^+^ (**A**) CD23^+^ and (**B**) BCL2^+^ GC B cells (left), BCL6^-^CD80^+^ GC transit B cells (middle), and MBCs (right). Data are combined from three independent experiments with 5-6 mice per group. (**C**) WT mice were infected with *P.ch* and treated with PBS or IL4C both with or without emricasan on days 12 and 14. Representative flow plots showing gating on MSP1^+^ B cells from mice treated with IL4C (top) or IL4C and emricasan (bottom) on day 15 post-infection. (**D**) Number of MSP1^+^ total B cells, GC B cells, and swIg^+^CD73^+^CD80^+^ MBCs in each group. (**E**) Flow plots and quantification of BCL6^-^ GC B cells. (**F**) Number of CD80^+^ GC transit B cells in each group. Data are combined from three independent experiments with 6 mice per group. See also Figure S5.

To test if the diminished expression of BCL2 in GC B cells following IL4C treatment is indicative of increased death of IL-4 “selected” pre-memory GC B cells and recently formed MBCs, we administered IL4C or PBS simultaneously with the pan-caspase inhibitor emricasan to WT mice infected with *P.ch* (**Figure 4C**). Emricasan treatment resulted in an increase in MSP1-specific B cells regardless of PBS or IL4C treatment, demonstrating that, as expected, blocking cell death increases MSP1-specific B cell numbers in the presence or absence of IL-4 treatment (**Figure 4D**). However, while mice treated with both PBS and emricasan had only modest increases in GC B cells and swIg^+^CD73^+^CD80^+^ MBCs compared to PBS-treated mice (fold changes of 1.22 and 0.15, respectively), mice receiving IL4C and emricasan had a <u>5-fold</u> increase in GC B cells and a <u>6-fold</u> increase in swIg^+^CD73^+^CD80^+^ MBCs compared to mice treated with IL4C alone, demonstrating that IL-4 is leading to increased GC B cell death (**Figure 4D**). Consistent with this finding, mice treated with IL4C and emricasan had a significantly higher number of BCL6^-^ GC B cells compared to mice treated with PBS and emricasan, indicating that there is a large proportion of GC B cells dying due to IL-4-mediated downregulation of BCL6 (**Figure 4E**). IL4C and emricasan treatment also resulted in an increase in CD80^+^ GC transit B cells compared to IL4C alone, suggesting that IL-4-mediated cell death is preventing pre-memory cells from surviving after exiting the GC (**Figure 4F**).

We additionally validated these results in BCL2 transgenic mice (BCL2-Tg), which have a functional human BCL2 transgene expressed only in the murine B cell lineage^10^. We infected WT and BCL2-Tg littermates with *P.ch* and treated them with IL4C or PBS on days 12 and 14 (**Figure S5C**). As with the use of emricasan, there was no loss in antigen-specific B cells between BCL2-Tg mice treated with PBS or IL-4 at day 15 (**Figure S5D**). Consistent with BCL2 acting antagonistically to the GC transcriptional program, BCL2-Tg mice did not form significantly larger GCs (**Figure S5D**). However, IL4C treatment in BCL2-Tg mice resulted in a significant increase in swIg^+^CD73^+^CD80^+^ MBCs compared to littermate controls, whereas PBS treatment in BCL2-Tg mice did not specifically impact swIg^+^CD73^+^CD80^+^ MBCs (**Figure S5D**). Together, these data demonstrate that the loss of B cells in response to IL4C treatment is due to the death of pre-memory swIg^+^ GC B cells following the IL-4-mediated downregulation of BCL6. Preventing this cell death through either caspase inhibition or overexpression of an anti-apoptotic signal fully recovers both GC B cells and GC-derived MBCs.

### IL-4 signaling regulates MBC selection stringency

Together, our data suggested that GC B cell-intrinsic IL-4 signaling has the capacity to drive the exit of nascent, CD23^+^CD80^+^ memory precursor cells from the GC. This raises the possibility that IL-4 from GC Tfh cells could be a key cue involved in the selection of somatically hypermutated (SHM) GC B cells. Germinal center selection is necessary to ensure that a diverse population of B cells with mutated BCRs of sufficient affinity survive into the MBC pool, and this selection process occurs via interactions with GC Tfh cells and antigen-bearing follicular dendritic cells^50^. The specific cues that direct this selection process remain unknown. Based on our data, we hypothesized that IL-4 from a GC Tfh cell can select B cells to exit the GC via the downregulation of BCL6. Additional cues to the B cell would lead to the expression of anti-apoptotic molecules like BCL2, which would then promote long-lived MBC survival. We therefore aimed to assess if modulating IL-4 signaling could alter GC B cell selection into the MBC pool. We hypothesized that the addition of exogenous IL4C would inappropriately “select” GC B cells that had not undergone sufficient cycles of somatic hypermutation and affinity maturation to have been selected under physiological conditions. To test this, we sorted MSP1-specific IgG^+^CD80^+^ GC transit B cells and IgG^+^CD73^+^CD80^+^ MBCs on day 15 following PBS or IL4C treatment on days 12 and 14 and bulk-sequenced their B cell receptors (BCRs). Cells from the IL4C-treatment group in both populations had fewer somatic hypermutations than those from control-treated mice, which has been shown to correlate with affinity^51^ (**Figure 5A**).

**Figure 5.**
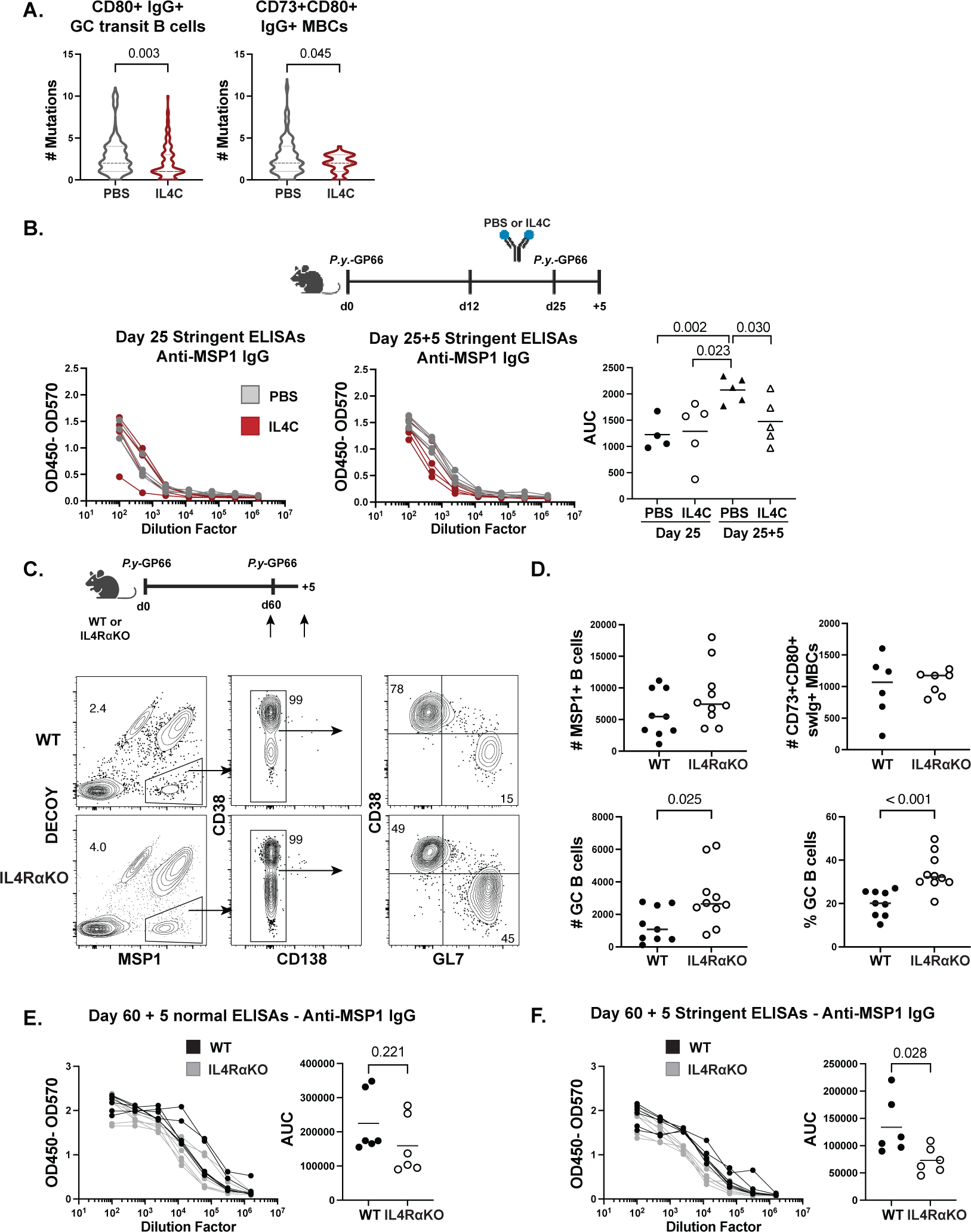
IL-4 signaling regulates MBC selection stringency. (**A**) Number of somatic hypermutations in MSP1^+^IgG^+^ CD80^+^ GC transit and CD73^+^CD80^+^ memory B cells sorted and sequenced on day 15 following PBS or IL4C treatments on days 12 and 14. Data are representative of two independent experiments with 6 mice per group. (**B**) WT mice were infected with *P.y.*-GP66 and treated with PBS or IL4C every other day between days 12-25, and re-challenged on day 25. Stringent ELISA dilution curves and AUC for anti-MSP1 IgG in serum collected at day 25 and 5 days post-challenge. Data are combined from two independent experiments with 4-5 mice per group. (**C**) WT and IL4RαKO mice were infected with *P.y*-GP66 and the MSP1-specific B cell response was analyzed on day 60, or mice were re-challenged on day 60 and analyzed 5 days post-challenge. Representative flow plots showing gating on MSP1^+^ B cells from each group. (**D**) Number of MSP1^+^ B cells and swIg^+^CD73^+^CD80^+^ MBCs (top) and number and frequency of GC B cells (bottom) at day 60. Data are combined from three independent experiments with 6-10 mice per group. (**E**) Normal and (**F**) stringent ELISA dilution curves and AUC for anti-MSP1 IgG in serum collected 5 days post-challenge. Data are combined from two independent experiments with 6 mice per group. See also Figure S6.

To verify that MBCs formed in the presence of excess IL-4 are in fact of lower affinity, we used a re-challenge model to induce memory B cell differentiation into plasmablasts, which allows for the analysis of MBC-derived serum antibodies by ELISA. We infected mice with *P.y.*-GP66 and administered PBS or IL4C every other day from day 12 through day 25, then re-challenged the mice with 10^7^ *P.y.*-GP66-iRBCs. We analyzed anti-MSP1 IgG antibody titers in the serum by ELISA under normal and chaotropic or “stringent” conditions using a urea wash to disrupt weak bonds to get a measure of antibody affinity/avidity. Serum was collected at both 25 days post-infection and 5 days post-challenge when swIg^+^ MBCs differentiate into plasmablasts^23^ (**Figure 5B**). There were no significant differences in serum anti-MSP1 IgG levels prior to challenge at day 25 as measured by normal or stringent ELISA suggesting that antibody quantity or quality present at this time point was not impacted by IL4C treatment ^52^ (**Figure 5B, Figure S6A**). After challenge, normal ELISAs revealed higher serum antibody titers in both challenged groups, although not significantly so for the IL4C treated animals. Furthermore, although there are not significant differences in antibody titers seen in stringent ELISAs on serum from the two groups at day 25, after rechallenge, stringent ELISAs showed a significant drop in signal from the IL4C-treated mice compared to the PBS treated mice (**Figure 5B**). This indicates that while early antibody secreting cells are not impaired in the IL4C treated mice after a primary infection, MBCs formed in the presence of excess IL-4 were of lower affinity than those formed in control-treated mice. Together, these results suggest that IL-4-mediated downregulation of BCL6 and the resulting increase in cell death underlies both the loss of MBCs and of high affinity MBCs.

These results led us to hypothesize that in an environment with physiological levels of IL-4 signaling, a lack of IL4-signaling may maintain longer GCs and produce lower affinity MBCs. To test this, we infected WT mice and global IL4Rα-deficient (IL4RαKO) mice with *P.y*-GP66 and analyzed the MSP1-specific B cell response at day 60 to determine if there were differences in GC maintenance or memory B cells (**Figure 5C**). Importantly, IL4RαKO mice have been shown to develop GCs at a similar rate to WT mice, likely due to the outsize role of IL-21 in promoting GC formation and in keeping with our data demonstrating equivalent BCL6 expression in STAT6KO B cells^15,53^. WT and IL4RαKO mice infected with *P.y*-GP66 had similar numbers of total MSP1-specific B cells and no differences in the number of swIg^+^CD73^+^CD80^+^ MBCs, demonstrating that IL-4 is not the only cue that can promote MBC formation (**Figure 5D**). However, IL4RαKO mice had significantly larger GCs at day 60 by both number and frequency (**Figure 5D**). The larger GCs cannot be attributed to a lack of parasite clearance, as the IL4RαKO mice controlled parasitemia as well as the WT mice throughout infection (**Figure S6B and S6C**). Together, these data suggest that while MBCs can still form, there may be increased retention or survival of GC B cells in the absence of IL-4 signaling. Importantly, there were no significant differences in serum anti-MSP1 IgG levels as measured by ELISA or by stringent ELISA (**Figure S6D and S6E**). Thus, while IL-4 may impact MBC survival and selection within the GC, it may not affect plasma cell selection.

We next sought to determine whether the longer lasting GCs were due to enhanced survival of lower affinity GC B cells, which would then result in the formation of lower affinity MBCs. To test this, we re-challenged WT and IL4RαKO mice with 10^7^ *P.y-*GP66-iRBCs at day 60 and analyzed the serum antibody response 5 days post-challenge, when MBCs have begun to differentiate into plasmablasts^23^ (**Figure 5C**). Serum anti-MSP1 IgG levels were similar between the two re-challenged groups by normal ELISA, indicating that the lack of IL-4 signaling does not impact MBC differentiation into plasmablasts (**Figure 5E**). Stringent ELISAs, however, revealed a significant reduction in signal in serum from IL4RαKO mice compared to serum from WT mice at day 5 post-challenge, suggesting that the bulk of IL4RαKO MBC-derived antibodies were lower affinity than those from WT mice (**Figure 5F**). This indicates that in the absence of IL-4-mediated downregulation of the anti-apoptotic BCL6, lower affinity B cells can persist in the GC, thus leading to a lower affinity MBC pool. Together, these findings suggest that IL-4 tightly regulates MBC selection in the GC, and that both too little or too much IL-4 signaling can lead to a loss of MBC selection stringency.

### A lack of IL-4 signaling in B cells promotes longer-lasting GCs and lower affinity MBCs

Although GC formation has shown to be largely normal in IL4RαKO mice, the lack of IL-4 signaling in other cell populations may be involved in the increased GC longevity and lower affinity MBCs observed above. To address this, we infected S1PR2creERT2^+/−^ Tdtomato^flox^IL4Rα^flox^ mice and their S1PR2creERT2^-/-^ littermates with *P.y.*-GP66 and fed them tamoxifen chow from day 10 through day 60 (**Figure 6A**). This model allows us to determine whether a specific lack of IL-4 signaling in GC B cells impacts GC persistence and MBC formation. As observed with these mice at day 20 (Figure 3A), there was an incomplete deletion of IL4Rα on TdTomato^+^ B cells, limiting numerical analyses between groups. Within the MSP1-specific TdTomato^+^ (S1PR2 fate-mapped) population from S1PR2creERT^+/−^ mice, there is no difference in the frequency of IL4Rα^-^ cells found within total B cells or swIg^+^CD73^+^CD80^+^ MBCs, indicating that GC-derived MBCs can still form in the absence of IL-4 signaling, as shown above (**Figure 6B**). However, there was a significantly increased proportion of IL4Rα^-^ B cells within the GC compared to other B cell subsets, suggesting these cells are more likely to be retained in the GC (**Figure 6B**). Similarly, there was a consistently higher frequency of GC B cells within the IL4Rα^-^TdTomato^+^ population compared to GC B cells within the IL4Rα^+^TdTomato^+^ population (**Figure 6C**). This recapitulates the findings in the IL4RαKO mice, demonstrating that GCs can persist for longer in the absence of the IL-4-mediated downregulation of BCL6, potentially due to increased GC B cell survival. Importantly, we also noted that within the MSP1-specific TdTomato^+^ population from S1PR2creERT^+/−^ mice, the tetramer mean fluorescence intensity (MFI) was significantly higher on IL4Rα^+^ TdTomato^+^swIg^+^CD73^+^CD80^+^ MBCs compared to their IL4Rα^-^ counterparts within the same animal (**Figure 6D**). The MFI of antigen binding has been shown to correlate with BCR affinity, thus this indicates that the absence of IL-4 signaling in GC B cells may result in the formation of lower affinity GC-derived MBCs^54–57^. This agrees with our observations in the IL4RαKO mice that the IL-4-mediated downregulation of BCL6 in GC B cells is necessary to limit the persistence of lower affinity GC B cells in the memory B cell pool.

**Figure 6.**
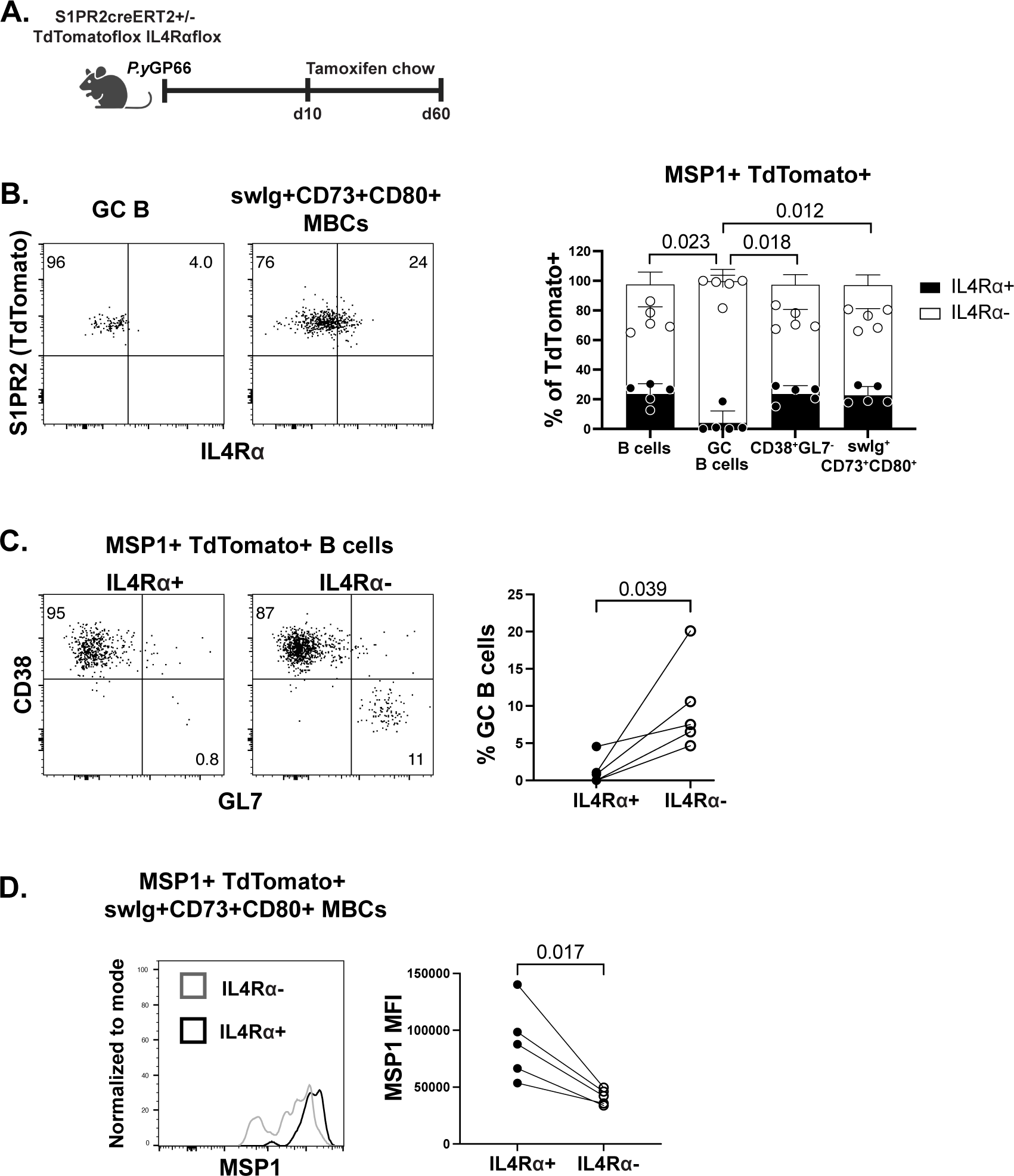
Lower affinity B cells persist into the memory pool in the absence of B cell-intrinsic IL-4 signaling. (**A**) S1PR2creERT2^+/−^TdTomato^flox^IL4Rα^flox^ mice and S1PR2creERT2^-/-^ littermates were infected with *P.y-*GP66 and fed tamoxifen chow beginning 10 days post-infection and analyzed on day 60. (**B**) Representative flow plots and quantification of IL4Rα^+^ and IL4Rα^-^ TdTomato^+^ B cells. (**C**) Representative flow plots and quantification of GC B cells within IL4Rα^+^ or IL4Rα^-^ MSP1-specific TdTomato^+^ B cells. (**D**) Representative histograms and quantification of MSP1 tetramer MFI on IL4Rα^+^ and IL4Rα^-^ MSP1^+^TdTomato^+^swIg^+^CD73^+^CD80^+^ MBCs. Data are combined from three independent experiments with 5 mice per group.

## Discussion

Our data support a model in which IL-4 can promote either GC B cell entry or exit depending on the amount of BCL6 expression in a given cell. Cells that express no to low levels of BCL6 enhance BCL6 expression in response to IL-4 signaling, while those that have high levels of BCL6, like GC B cells, downregulate BCL6 in response to IL-4. More specifically, as previously shown, IL-4 produced by NKT cells or Tfh cells early in an immune response acts upon naive and early activated B cells to promote BCL6 expression and thereby promote migration into GCs^17,18,58^. In multiple models, there continues to be a significant amount of IL-4 produced by CD4^+^ T cells following GC formation within the GC itself^59–61^, consistent with our own findings that GC Tfh are the predominant CD4^+^ T cell source of IL-4 in response to *Plasmodium* infection. However, the role of IL-4 within the GC has only recently been examined through our work and very recently from the work of Duan and colleagues^16^, which demonstrated that unrestricted IL-4 availability in the GC can constrain memory cell formation. In keeping with earlier reports that IL-4 is broadcast around a T cell instead of secreted directly at the synapse^62^, their data suggested that IL4Rα expressed on FDCs serves as a sink for excess IL-4, thereby limiting its effects on bystander GC B cells that are not actively engaged with a GC Tfh cell. Importantly, these studies did not define the mechanism via which IL-4 was acting on GC B cells.

Herein, we demonstrate that IL-4 acts on GC B cells to downregulate BCL6, whether administered exogenously or perceived *in situ*, by enhancing BCL6 expression to a threshold level that triggers its own negative autoregulation^19,41,42^. This correlates with increased cell death, which we found to be associated with a loss in selection stringency as measured by both a decrease in somatic hypermutation as well as in the affinity of antibodies from MBC-derived plasmablasts. We found that in the absence of IL4Rα signaling, B cells were preferentially able to remain in the GC, although they could still exit to form MBCs. This suggested a potential lack of the positive selection that would limit survival to high affinity clones over their lower affinity counterparts. Indeed, we determined that a global or GC B cell-intrinsic loss of IL-4 signaling resulted in the formation of lower affinity MBCs. Together, our findings indicate that BCL6 serves as a tunable switch responding to T cell IL-4 secretion, thereby releasing cells that have acquired adequate help and have therefore been selected.

We additionally identified a novel approach to distinguish memory precursors that have exited the GC through a combination of differing BCL6 and CD80 expression. This allowed us to determine that while IL-4 increased the proportion of exiting GC B cells, it also increased GC B cell death, as evidenced by the higher number of BCL6^-^ GC B cells and CD80^+^ GC transit B cells found when mice were treated with IL4C in conjunction with the pan-caspase inhibitor emricasan. One possible explanation for the IL-4-mediated cell death was the finding that GC B cells did not appropriately upregulate the anti-apoptotic protein BCL2 in response to IL4C treatment. It has previously been suggested that during a selection event, the downregulation of BCL6 should release repression of BCL2, as BCL2 is required for proper MBC survival^33,49^. Our findings therefore decouple these two events, and raise the possibility that two distinct signals from T cells are required for MBC selection: IL-4 to downregulate BCL6 and release cells from the GC transcriptional program, and an additional signal to upregulate MBC survival signals. It is likely that the loss of MBCs in response to increased IL-4 observed in our work and that of Duan and colleagues is due to an imbalance in the number of GC B cells that are downregulating BCL6 and the number of the GC B cells that can receive additional T cell or BCR survival signals^16^. While the additional required signal(s) is still unknown, previous studies have dissected how the strength of T cell-B cell receptor-ligand interactions can dictate whether a GC B cell migrates to the DZ or differentiates into a memory B cell or plasma cell, thus the amount of IL-4 received may also factor into these fate choices^2,5,39,63^.

Interestingly, our results indicated that IL-4 impacts MBC selection, but did not appear to impact plasma cell selection. Similarly, IL-21 has been shown to play an important role in clonal expansion in the DZ as well as in long-lived plasma cell formation, but has not been shown to significantly impact MBC formation^46,64–66^. What makes a Tfh cell secrete IL-4 versus IL-21 in response to an interaction with a GC B cell has not been determined, however it is known that very few Tfh cells can produce both cytokines simultaneously^26^. IL-4 and IL-21 production appear to be relatively segregated to the light zone or dark zone, respectively^15,26,61^, suggesting cytokine signals in the GC may be temporally and spatially regulated, and perhaps memory B cell versus plasma cell fate are as well.

The signals underlying GC B cell selection and memory B cell exit have remained elusive, in part due to the complex nature of cellular interactions in the GC. The finding that IL-4 can promote GC entry early in an immune response as well as initiate selection and exit later in the GC illuminates a significant role for cytokines in tuning BCL6, which may go beyond the GC. How GC B cells integrate IL-4 with other cytokine or receptor-ligand signals remains to be determined, yet the ability to modulate IL-4 throughout an immune response may on its own allow for the manipulation of MBC affinity in vaccine settings.

## Acknowledgements

The authors thank the Pepper lab members as well as the Oberst and Gerner labs for helpful discussions and technical assistance. We are incredibly appreciative of Nate Bloom, Brian Freeman, Sonya Haupt, and Dr. Shane Crotty for providing us with tissues from the ΔBPS1 mice. We would also like to thank Dr. Chris Allen for the BCL2-Tg mice, and the University of Washington Department of Immunology Flow Cytometry Core and vivarium for their support. Experiment schematics in figures were created with <u>BioRender.com</u>. This work was supported by NIH T32 AI206677 and the National Science Foundation Graduate Research Fellowship Program under Grant No. DGE-1762114 (L.S), as well as NIH R01 AI118803, U01 AI42001, and BWF 1016766 (M.P.).

## Author contributions

L.S. and M.P. conceived and designed the study. L.S., C.D.T., and L.P. performed experiments. L.S. and C.D.T. analyzed data. B.H. provided technical and conceptual expertise. D.R. and J.C. provided reagents. L.S. and M.P. wrote the manuscript. C.D.T., B.H., L.P., D.R., and J.C. read and edited the manuscript.

## Methods

### Resource and Materials Availability

Further information and requests for resources and reagents should be directed to and will be fulfilled by the lead contact, Marion Pepper.

### Mice

C57BL/6, CD45.1^+^ (B6.SJL-*Ptprc^a^ Pepc^b^/BoyJ*), and STAT6KO (B6.129S2(C)-*Stat6^tm1Gru^*/J)^67^ mice were purchased from Jackson Laboratories. S1PR2creERT2^+/−^TdTomato^flox^ (Tg(S1PR2-cre/ERT2)) mice were provided by Dr. Tomohiro Kurosaki^5^. This strain was crossed to IL4Ra^flox^ (IL4ra^tm2Fbb^) mice^68^, provided by Dr. Jakob von Moltke, to generate S1PR2creERT2^+/−^ TdTomato^flox^IL4Ra^flox^ mice. IL4RαKO (IL-4Rα^-/-^) mice were provided by Dr. Frank Brombacher and were bred and maintained in our laboratory^69^. KN2^+/+^ mice were provided by Dr. Markus Mohrs^70^. This strain was crossed to C56BL/6 mice to generate KN2^+/−^ mice. ΔBPS1 mice were provided by Dr. Jinyong Choi^19^. BCL2-Tg (B6.Cg-Tg(BCL2)22Wehi/J) mice were provided by Dr. Chris Allen^71^. All mice were bred and housed under specific-pathogen free conditions at the University of Washington. All experiments were performed in accordance with the University of Washington Institutional Care and Use Committee guidelines.

### Infections and parasitemia analysis

*Plasmodium chabaudi chabaudi (AS)* parasites and *Plasmodium yoelii 17XNL-GP66* parasites were maintained as frozen blood stocks and passaged through donor mice. Primary mouse infections were initiated by intraperitoneal (i.p.) injection of 10^6^ infected red blood cells (iRBCs) from donor mice. Re-challenge experiments were performed by injecting mice with 10^7^ iRBCs i.p. 60 days after primary injection. Parasitemia was measured by flow cytometry by fixing a drop of blood with 0.025% glutaraldehyde, then staining with CD45 APC (Clone 30-F11, BD Biosciences), Ter119 FITC (BD Biosciences), CD71 PE (Clone R17217, Invitrogen), and Hoechst33342. For ΔBPS1 *in vitro* experiments, mice were infected with 2×10^5^ PFU LCMV_arm_ i.p.

### Tetramers

The GP66_-77_:I-A^b^-APC tetramer (I-A^b^/LCMV.GP66.DIYKGVYQFKSV) used for isolating GP66-specific T cells was obtained from the National Institutes of Health Tetramer Core. For *P.ch* MSP1-specific B cell isolation, recombinant His-tagged C-terminal MSP1 protein (amino acids 4960 to 5301) from *P.ch* was produced by *Pichia pastoris* and purified using a Ni-NTA agarose column as previously described^72^. For *P.y*-GP66 MSP1-specific B cell isolation, a plasmid containing His-tagged 14 kD truncated carboxy terminus of *P.y* MSP1 protein (amino acids 1662-1757) was transfected into HEK293F cells and purified using a Ni-NTA agarose column as described previously^73^. Purified *P.ch* MSP1 protein and purified *P.y* MSP1 protein were biotinylated using the EZ-Link Sulfo-NHS-LC Biotinylation Kit (ThermoFisher) and tetramerized with streptavidin-PE (Agilent) or streptavidin-APC (Agilent) as previously described^23,74^. The PE decoy reagent to gate out non-MSP1-specific B cells was generated by conjugating SA-PE to AF647 using an AF647 antibody labeling kit (ThermoFisher Scientific), washing and removing any unbound AF647, and incubating with an excess of irrelevant biotinylated HIS-tagged protein. The APC decoy reagent was generated in the same manner, by conjugating SA-APC to Dylight 755 using a DyLight 755 antibody labeling kit (ThermoFisher Scientific).

### Injections

For the administration of complexed IL-4 (IL4C), 5 ug of recombinant mouse IL-4 (BioLegend) was incubated with 25 ug of anti-mouse IL-4 antibody (11B11; BioXCell) for 2 minutes at room temperature (RT). The IL4C was then diluted to 150 uL with sterile PBS and administered intravenously (i.v.). For the administration of non-complexed IL-4, 5 ug of recombinant mouse IL-4 (BioLegend) was diluted to 150 uL with sterile PBS and administered i.v. Emricasan (Sigma-Aldrich) was resuspended to 2mg/mL in DMSO, prepared at 2.5 mg/kg in pre-warmed sterile PBS, and injected i.p.

### Mouse cell enrichment and flow cytometry

Splenic cell suspensions were prepared and filtered through 100 uM Nitex mesh (Amazon). For B cell enrichment, cells were resuspended in 200 uL PBS containing 2% FBS and Fc block (2.4G2) and incubated with decoy tetramer at a concentration of 10 nM for 10 min at RT. MSP1-PE or -APC tetramer was added at a concentration of 10 nM and cells were incubated for 20 min on ice. Cells were washed, incubated with anti-PE or anti-APC magnetic beads (Miltenyi) for 30 min on ice, and passed over magnetized LS columns (Miltenyi) to elute the bound cells as previously described^74^. For T cell enrichment, cells were resuspended in 200uL PBS containing 2% FBS and incubated with 10 nM GP66-77:I-A^b^-APC tetramer for 1 hour at RT. Cells were washed, incubated with anti-APC magnetic beads, and passed over magnetized LS columns. All bound cells were stained with surface antibodies followed by fixation and intracellular antibody staining when needed (**Table S1**). Cell counts were determined using Accucheck cell counting beads. Cells were analyzed on the Cytek Aurora and analyzed using FlowJo 10 software (Treestar).

### *In vitro* splenocyte cultures

Splenic cell suspensions were prepared and filtered through 100 uM Nitex mesh. For negative selection of naive B cells, splenocytes were resuspended in 200 uL PBS containing 2% FBS and Fc block and incubated with biotin anti-mouse F4/80 (Clone BM8, BioLegend), CD3 (Clone 500A2, BD Biosciences), CD4 (Clone RM4-5, eBioscience), CD8 (Clone 53-6.7, eBioscience), CD11b (Clone M1/70, eBioscience), CD11c (Clone N418, eBioscience), CD43 (Clone eBioR2/60, Invitrogen), Ly6G (Clone RP6-8C5, eBioscience), Ter119 (Clone TER-119, eBioscience), and CD138 (Clone 281-2, BD Biosciences) for 20 min on ice. For negative selection of GL7^+^ B cells, cells were additionally incubated with biotin anti-mouse CD38 (Clone 90/CD38, BD Biosciences), IgD (Clone 11-26c, Invitrogen), and CD49b (Clone DX5, BD Biosciences). In both cases, cells were washed, incubated with anti-biotin magnetic beads (Miltenyi) for 30 min on ice, and passed over magnetized LS columns. Flow through was collected and an aliquot of cells were checked for purity prior to culture. 1 million B cells/well were added to 12-well plates in 1 mL DMEM (Gibco) supplemented with 10% FBS (R&D Systems), 1% penicillin/streptomycin (ThermoFisher Scientific), 15 mM Hepes (Gibco), and 55 uM β-mercaptoethanol (Gibco). 1 ug/mL of agonistic anti-CD40 antibody (Clone FGK4.5, BioXCell) diluted in PBS was added to all wells, and 10 ng/mL of recombinant mouse IL-4 (BioLegend) diluted in PBS was added to wells cultured with the cytokine. Plates were incubated at 37°C in a 5% CO2 incubator for 24h, then cells were harvested and washed with PBS containing 2% FBS and stained for flow cytometry as described above.

### Adoptive cell transfers

Single cell suspensions from ΔBPS1 spleens were prepared and filtered through 100uM Nitex mesh. Red blood cells were lysed with ACK buffer (ThermoFisher Scientific). Naive B cells were isolated by negative magnetic activated cell sorting using the EasySep^TM^ Mouse B cell Isolation Kit (StemCell Technologies). 30 million B cells were injected i.p. into CD45.1^+^ mice.

### ELISAs

96-well plates (Corning) were coated with 2 ug/mL of recombinant *P.ch* or *P.y* MSP1 diluted in PBS and incubated at 4°C overnight. Plates were washed with PBS containing 0.05% Tween-20 (PBS-T) and incubated with blocking buffer (PBS-T and 3% milk) for 1 hour at RT. Serum was serially diluted in dilution buffer (PBS-T and 1% milk), added to plates, and incubated at RT for 2 hours. Plates were washed and incubated with biotin anti-mouse IgG (BioLegend) for 1 hour at RT, followed by Streptavidin-HRP (BD) for 30 min at RT. Bound antibodies were detected with 1X TMB (Invitrogen) and quenched with 1M HCl. Sample optical density (OD) was measured at 450 nM and 570 nM using an iMark Microplate Reader (Bio-Rad).

### ELISpots

96-well ELISpot plates (Millipore) were coated with 20 ug/mL of recombinant *P.ch* or *P.y* MSP1 diluted in PBS and incubated at 4°C overnight. Plates were washed with PBS-T and blocked with 1% BSA and 5% sucrose in PBS for 2h at RT. Single cell suspensions of murine bone marrow were prepared and passed over 100 uM Nitex mesh. Following red blood cell lysis, cells were plated onto coated ELIspot plates and incubated at 37°C for 5h. Cells were washed off and antibody secreting cells were detected using IgG biotin (BioLegend) followed by streptavidin-alkalkine phosphatase (R&D Systems). Spots were developed using BCIP/NBT (Mabtech) and were counted and analyzed using the CTL ELIspot reader and Immunospot analysis software (Cellular Technology Limited). Non-specific and background spots were determined by wells containing no cells. The number of spots detected per well was used to calculate the spot frequency per 10,000 cells.

### B cell sorting and BCR sequencing

Cells were processed and stained as described above, and sorted on a FACS Aria II cell sorter (BD) into SMART-Seq lysis buffer (Takara). cDNA was generated for each sample using the SMART-Seq v4 kit (Takara). Samples were end-repaired, A-tailed, and ligated with custom UMI adapters (IDT) (**Table S2**) at 2 uM according to NEBNext Ultra II instructions (NEB). Libraries were prepped following NEBNext Immune Sequencing Kit’s protocol (NEB), and qPCR was used to determine cycling conditions prior to plateau. Samples were pooled at equimolar concentration and ran on a MiSeq (Illumina) with a MiSeq Reagent Kit v3 (600-cycle) (Illumina).

### BCR sequence processing

All samples were processed with an in-house pipeline using pRESTO, BBDuk, and custom python scripts^75,76^. Sequences were filtered for Q20 average quality and the TSO sequence. Consensus sequences were made using the UMI, and leader sequences were removed. Paired ends were assembled and sequences with bases below Q30 were removed. Sequences were collapsed to remove duplicates and all sequences filtered for at least 2 reads. Resulting fastq files were submitted to IMGT for alignment. Clones were identified using Shazam and Change-O with a clustering threshold of 0.04^77^. Sequences were analyzed further in R (4.0.4) using tidyverse (1.3.0)^78^.

### Histology and image analysis

Spleens were isolated from infected mice and immediately submerged in BD Cytofix diluted 1:3 with PBS for 24h at 4°C. Spleens were then washed with PBS and dehydrated in 30% sucrose for 24h at 4°C before being embedded in OCT freezing medium. 18 uM sections were cut on a cryostat and stained with anti-mouse IgD ef450 (Clone 11-26c, Invitrogen), CD3 AF488 (Clone 17A2, BioLegend), BCL6 AF647 (Clone K112-91, BD Biosciences), and CD35 biotin (Clone 8C12, BD Biosciences), followed by Streptavidin Cy3 (Jackson ImmunoResearch). Slides were cover-slipped with Fluoromount G mounting media (SouthernBiotech) and images were acquired on a Nikon Eclipse 90i with NIS-Elements software. Imaris (Bitplane) was used for image analysis. GC diameter and circumference were measured as the area around or across the BCL6^+^ cells using Imaris, with each measurement taken three times and averaged.

### Statistical analysis

All statistical analyses were performed in GraphPad Prism 9 (GraphPad Software). Unpaired, two-tailed Student’s t tests were applied to determine the differences between two individual groups. Paired, two-tailed Student’s t tests were applied to determine the differences between two groups in adoptive transfer experiments. In the event of three or four groups, a one-way ANOVA followed by Tukey’s multiple comparisons test was performed. The statistical details of each experiment are indicated in the figure legends.

## References

1. Mesin, L., Ersching, J., and Victora, G.D. (2016). Germinal Center B Cell Dynamics. Immunity 45, 471–482. 10.1016/j.immuni.2016.09.001.

2. Ise, W., Fujii, K., Shiroguchi, K., Ito, A., Kometani, K., Takeda, K., Kawakami, E., Yamashita, K., Suzuki, K., Okada, T., et al. (2018). T Follicular Helper Cell-Germinal Center B Cell Interaction Strength Regulates Entry into Plasma Cell or Recycling Germinal Center Cell Fate. Immunity 48, 702–715.e4. 10.1016/J.IMMUNI.2018.03.027.

3. Inoue, T., Moran, I., Shinnakasu, R., Phan, T.G., and Kurosaki, T. (2018). Generation of memory B cells and their reactivation. Immunol Rev 283, 138–149. 10.1111/imr.12640.

4. Inoue, T., Shinnakasu, R., Kawai, C., Ise, W., Kawakami, E., Sax, N., Oki, T., Kitamura, T., Yamashita, K., Fukuyama, H., et al. (2021). Exit from germinal center to become quiescent memory B cells depends on metabolic reprograming and provision of a survival signal. J Exp Med 218. 10.1084/jem.20200866.

5. Shinnakasu, R., Inoue, T., Kometani, K., Moriyama, S., Adachi, Y., Nakayama, M., Takahashi, Y., Fukuyama, H., Okada, T., and Kurosaki, T. (2016). Regulated selection of germinal-center cells into the memory B cell compartment. Nat Immunol 17, 861–869. 10.1038/ni.3460.

6. Haberman, A.M., Gonzalez, D.G., Wong, P., Zhang, T., and Kerfoot, S.M. (2019). Germinal center B cell initiation, GC maturation, and the coevolution of its stromal cell niches. Immunol Rev 288, 10–27. 10.1111/imr.12731.

7. Ochiai, K., Maienschein-Cline, M., Simonetti, G., Chen, J., Rosenthal, R., Brink, R., Chong, A.S., Klein, U., Dinner, A.R., Singh, H., et al. (2013). Transcriptional Regulation of Germinal Center B and Plasma Cell Fates by Dynamical Control of IRF4. Immunity 38, 918–929. 10.1016/j.immuni.2013.04.009.

8. Yeh, C.H., Nojima, T., Kuraoka, M., and Kelsoe, G. (2018). Germinal center entry not selection of B cells is controlled by peptide-MHCII complex density. Nat Commun 9. 10.1038/S41467-018-03382-X.

9. Jacobsen, J.T., Hu, W., Castro, T.B.R., Solem, S., Galante, A., Lin, Z., Allon, S.J., Mesin, L., Bilate, A.M., Schiepers, A., et al. (2021). Expression of Foxp3 by T follicular helper cells in end-stage germinal centers. Science 373. 10.1126/SCIENCE.ABE5146.

10. Smith, K.G.C., Weiss, U., Rajewsky, K., Nossal, G.J.V., and Tarlinton, D.M. (1994). Bcl-2 increases memory B cell recruitment but does not perturb selection in germinal centers. Immunity 1, 803–813. 10.1016/S1074-7613(94)80022-7.

11. Dominguez-Sola, D., Victora, G.D., Ying, C.Y., Phan, R.T., Saito, M., Nussenzweig, M.C., and Dalla-Favera, R. (2012). The proto-oncogene MYC is required for selection in the germinal center and cyclic reentry. Nat Immunol 13, 1083–1091. 10.1038/ni.2428.

12. Laidlaw, B.J., Duan, L., Xu, Y., Vazquez, S.E., and Cyster, J.G. (2020). The transcription factor Hhex cooperates with the corepressor Tle3 to promote memory B cell development. Nat Immunol 21, 1082–1093. 10.1038/s41590-020-0713-6.

13. Saito, M., Gao, J., Basso, K., Kitagawa, Y., Smith, P.M., Bhagat, G., Pernis, A., Pasqualucci, L., and Dalla-Favera, R. (2007). A Signaling Pathway Mediating Downregulation of BCL6 in Germinal Center B Cells Is Blocked by BCL6 Gene Alterations in B Cell Lymphoma. Cancer Cell 12, 280–292. 10.1016/j.ccr.2007.08.011.

14. Kuo, T.C., Shaffer, A.L., Haddad, J., Yong, S.C., Staudt, L.M., and Calame, K. (2007). Repression of BCL-6 is required for the formation of human memory B cells in vitro. J Exp Med 204, 819. 10.1084/JEM.20062104.

15. Gonzalez, D.G., Cote, C.M., Patel, J.R., Smith, C.B., Zhang, Y., Nickerson, K.M., Zhang, T., Kerfoot, S.M., and Haberman, A.M. (2018). Nonredundant Roles of IL-21 and IL-4 in the Phased Initiation of Germinal Center B Cells and Subsequent Self-Renewal Transitions. The Journal of Immunology 201, 3569–3579. 10.4049/JIMMUNOL.1500497.

16. Duan, L., Liu, D., An, J., Laidlaw, B.J., Correspondence, J.G.C., Chen, H., Mintz, M.A., Chou, M.Y., Kotov, D.I., Xu, Y., et al. (2021). Follicular dendritic cells restrict interleukin-4 availability in germinal centers and foster memory B cell generation. Immunity 54. 10.1016/j.immuni.2021.08.028.

17. Chevrier, S., Kratina, T., Emslie, D., Tarlinton, D.M., and Corcoran, L.M. (2017). IL4 and IL21 cooperate to induce the high Bcl6 protein level required for germinal center formation. Immunol Cell Biol 95, 925–932. 10.1038/icb.2017.71.

18. Gaya, M., Barral, P., Burbage, M., Aggarwal, S., Montaner, B., Warren Navia, A., Aid, M., Tsui, C., Maldonado, P., Nair, U., et al. (2018). Initiation of Antiviral B Cell Immunity Relies on Innate Signals from Spatially Positioned NKT Cells. Cell 172, 517–533.e20. 10.1016/J.CELL.2017.11.036.

19. Choi, J., Diao, H., Faliti, C.E., Truong, J., Rossi, M., Bélanger, S., Yu, B., Goldrath, A.W., Pipkin, M.E., and Crotty, S. (2020). Bcl-6 is the nexus transcription factor of T follicular helper cells via repressor-of-repressor circuits. Nat Immunol 21, 777–789. 10.1038/s41590-020-0706-5.

20. Ci, W., Polo, J.M., Cerchietti, L., Shaknovich, R., Wang, L., Shao, N.Y., Ye, K., Farinha, P., Horsman, D.E., Gascoyne, R.D., et al. (2009). The BCL6 transcriptional program features repression of multiple oncogenes in primary B cells and is deregulated in DLBCL. Blood 113, 5536–5548. 10.1182/BLOOD-2008-12-193037.

21. Kumagai, T., Miki, T., Kikuchi, M., Fukuda, T., Miyasaka, N., Kamiyama, R., and Hirosawa, S. (1999). The proto-oncogene Bcl6 inhibits apoptotic cell death in differentiation-induced mouse myogenic cells. Oncogene 18, 467–475. 10.1038/sj.onc.1202306.

22. Ranuncolo, S.M., Polo, J.M., Dierov, J., Singer, M., Kuo, T., Greally, J., Green, R., Carroll, M., and Melnick, A. (2007). Bcl-6 mediates the germinal center B cell phenotype and lymphomagenesis through transcriptional repression of the DNA-damage sensor ATR. Nat Immunol 8, 705–714. 10.1038/ni1478.

23. Krishnamurty, A.T., Thouvenel, C.D., Portugal, S., Keitany, G.J., Kim, K.S., Holder, A., Crompton, P.D., Rawlings, D.J., and Pepper, M. (2016). Somatically Hypermutated Plasmodium-Specific IgM(+) Memory B Cells Are Rapid, Plastic, Early Responders upon Malaria Rechallenge. Immunity 45, 402–414. 10.1016/j.immuni.2016.06.014.

24. Arroyo, E.N., and Pepper, M. (2020). B cells are sufficient to prime the dominant CD4+ Tfh response to Plasmodium infection. Journal of Experimental Medicine 217. 10.1084/JEM.20190849.

25. Langhorne, J., Gillard, S., Simon, B., Slade, S., and Eichmann, K. (1989). Frequencies of CD4+ T cells reactive with plasmodium chabaudi chabaudi: Distinct response kinetics for cells with Th1 and Th2 characteristics during infection. Int Immunol 1, 416–424. 10.1093/intimm/1.4.416.

26. Weinstein, J.S., Herman, E.I., Lainez, B., Licona-Limón, P., Esplugues, E., Flavell, R., and Craft, J. (2016). TFH cells progressively differentiate to regulate the germinal center response. Nat Immunol 17, 1197–1205. 10.1038/ni.3554.

27. Katona, I.M., Maliszewski F D Finkelman, C.R., Madden, K.B., Morris, S.C., and Holmes, J.M. (2021). cytokine-anti-cytokine antibody complexes. cytokines by injection of Prolongation of in vivo effects of exogenous Anti-cytokine antibodies as carrier proteins.

28. Taylor, J.J., Pape, K.A., and Jenkins, M.K. (2012). A germinal center-independent pathway generates unswitched memory B cells early in the primary response. J Exp Med 209, 597–606. 10.1084/jem.20111696.

29. Wang, Y., Shi, J., Yan, J., Xiao, Z., Hou, X., Lu, P., Hou, S., Mao, T., Liu, W., Ma, Y., et al. (2017). Germinal-center development of memory B cells driven by IL-9 from follicular helper T cells. Nat Immunol 18, 921–930. 10.1038/ni.3788.

30. Laidlaw, B.J., Schmidt, T.H., Green, J.A., Allen, C.D.C., Okada, T., and Cyster, J.G. (2017). The Eph-related tyrosine kinase ligand Ephrin-B1 marks germinal center and memory precursor B cells. J Exp Med 214, 639–649. 10.1084/jem.20161461.

31. Kerfoot, S.M., Yaari, G., Patel, J.R., Johnson, K.L., Gonzalez, D.G., Kleinstein, S.H., and Haberman, A.M. (2011). Germinal Center B Cell and T Follicular Helper Cell Development Initiates in the Interfollicular Zone. Immunity 34, 947–960. 10.1016/j.immuni.2011.03.024.

32. Huang, C., Gonzalez, D.G., Cote, C.M., Jiang, Y., Hatzi, K., Teater, M., Dai, K., Hla, T., Haberman, A.M., and Melnick, A. (2014). The BCL6 RD2 domain governs commitment of activated B cells to form germinal centers. Cell Rep 8, 1497–1508. 10.1016/J.CELREP.2014.07.059.

33. Laidlaw, B.J., and Cyster, J.G. (2021). Transcriptional regulation of memory B cell differentiation. Nat Rev Immunol 21, 209–220. 10.1038/s41577-020-00446-2.

34. Niu, H., Cattoretti, G., and Dalla-Favera, R. (2003). BCL6 controls the expression of the B7-1/CD80 costimulatory receptor in germinal center B cells. Journal of Experimental Medicine 198, 211–221. 10.1084/jem.20021395.

35. Tomayko, M.M., Steinel, N.C., Anderson, S.M., and Shlomchik, M.J. (2010). Cutting Edge: Hierarchy of Maturity of Murine Memory B Cell Subsets. The Journal of Immunology 185, 7146–7150. 10.4049/JIMMUNOL.1002163.

36. Zuccarino-Catania, G. V., Sadanand, S., Weisel, F.J., Tomayko, M.M., Meng, H., Kleinstein, S.H., Good-Jacobson, K.L., and Shlomchik, M.J. (2014). CD80 and PD-L2 define functionally distinct memory B cell subsets that are independent of antibody isotype. Nature Immunology 2014 15:7 15, 631–637. 10.1038/ni.2914.

37. Weisel, F.J., Mullett, S.J., Elsner, R.A., Menk, A. V., Trivedi, N., Luo, W., Wikenheiser, D., Hawse, W.F., Chikina, M., Smita, S., et al. (2020). Germinal center B cells selectively oxidize fatty acids for energy while conducting minimal glycolysis. Nat Immunol 21, 331. 10.1038/S41590-020-0598-4.

38. Luo, W., Conter, L., Elsner, R.A., Smita, S., Weisel, F., Callahan, D., Wu, S., Chikina, M., and Shlomchik, M. (2023). IL-21R signal reprogramming cooperates with CD40 and BCR signals to select and differentiate germinal center B cells. Sci Immunol 8. 10.1126/SCIIMMUNOL.ADD1823.

39. Good-Jacobson, K.L., Song, E., Anderson, S., Sharpe, A.H., and Shlomchik, M.J. (2012). CD80 Expression on B Cells Regulates Murine T Follicular Helper Development, Germinal Center B Cell Survival, and Plasma Cell Generation. The Journal of Immunology 188, 4217–4225. 10.4049/jimmunol.1102885.

40. Green, J.A., Suzuki, K., Cho, B., Willison, L.D., Palmer, D., Allen, C.D.C., Schmidt, T.H., Xu, Y., Proia, R.L., Coughlin, S.R., et al. (2011). The sphingosine 1-phosphate receptor S1P2 maintains germinal center B cell homeostasis and promotes niche confinement. Nat Immunol 12, 672. 10.1038/NI.2047.

41. Pasqualucci, L., Migliazza, A., Basso, K., Houldsworth, J., Chaganti, R.S.K., and Dalla-Favera, R. (2003). Mutations of the BCL6 proto-oncogene disrupt its negative autoregulation in diffuse large B-cell lymphoma. Blood 101, 2914–2923. 10.1182/blood-2002-11-3387.

42. Mendez, L.M., Polo, J.M., Yu, J.J., Krupski, M., Ding, B.B., Melnick, A., and Ye, B.H. (2008). CtBP Is an Essential Corepressor for BCL6 Autoregulation. Mol Cell Biol 28, 2175–2186. 10.1128/mcb.01400-07.

43. Schroder, A.J., Pavlidis, P., Arimura, A., Capece, D., and Rothman, P.B. (2002). Cutting Edge: STAT6 Serves as a Positive and Negative Regulator of Gene Expression in IL-4-Stimulated B Lymphocytes. The Journal of Immunology 168, 996–1000. 10.4049/jimmunol.168.3.996.

44. Wurster, A.L., Withers, D.J., Uchida, T., White, M.F., and Grusby, M.J. (2002). Stat6 and IRS-2 Cooperate in Interleukin 4 (IL-4)-Induced Proliferation and Differentiation but Are Dispensable for IL-4-Dependent Rescue from Apoptosis. Mol Cell Biol 22, 117. 10.1128/MCB.22.1.117-126.2002.

45. Linterman, M.A., Beaton, L., Yu, D., Ramiscal, R.R., Srivastava, M., Hogan, J.J., Verma, N.K., Smyth, M.J., Rigby, R.J., and Vinuesa, C.G. (2010). IL-21 acts directly on B cells to regulate Bcl-6 expression and germinal center responses. Journal of Experimental Medicine 207, 353–363. 10.1084/jem.20091738.

46. Zotos, D., Coquet, J.M., Zhang, Y., Light, A., D’Costa, K., Kallies, A., Corcoran, L.M., Godfrey, D.I., Toellner, K.-M., Smyth, M.J., et al. (2010). IL-21 regulates germinal center B cell differentiation and proliferation through a B cell-intrinsic mechanism. J Exp Med 207, 365–378. 10.1084/jem.20091777.

47. Nakagawa, R., Toboso-Navasa, A., Schips, M., Young, G., Bhaw-Rosun, L., Llorian-Sopena, M., Chakravarty, P., Sesay, A.K., Kassiotis, G., Meyer-Hermann, M., et al. (2021). Permissive selection followed by affinity-based proliferation of GC light zone B cells dictates cell fate and ensures clonal breadth. Proceedings of the National Academy of Sciences 118. 10.1073/PNAS.2016425118.

48. Smith, K.G.C., Light, A., O’Reilly, L.A., Ang, S.M., Strasser, A., and Tarlinton, D. (2000). bcl-2 Transgene expression inhibits apoptosis in the germinal center and reveals differences in the selection of memory B cells and bone marrow antibody-forming cells. Journal of Experimental Medicine 191, 475–484. 10.1084/jem.191.3.475.

49. Bhattacharya, D., Cheah, M.T., Franco, C.B., Hosen, N., Pin, C.L., Sha, W.C., and Weissman, I.L. (2007). Transcriptional Profiling of Antigen-Dependent Murine B Cell Differentiation and Memory Formation. The Journal of Immunology 179, 6808–6819. 10.4049/jimmunol.179.10.6808.

50. Victora, G.D., and Nussenzweig, M.C. (2022). Germinal Centers. 10.1146/annurev-immunol-120419-022408 40, 413–442. 10.1146/ANNUREV-IMMUNOL-120419-022408.

51. Weisel, F.J., Zuccarino-Catania, G.V., Chikina, M., and Shlomchik, M.J. (2016). A Temporal Switch in the Germinal Center Determines Differential Output of Memory B and Plasma Cells. Immunity 44, 116–130. 10.1016/J.IMMUNI.2015.12.004.

52. Tauzin, A., Gong, S.Y., Beaudoin-Bussières, G., Vézina, D., Gasser, R., Nault, L., Marchitto, L., Benlarbi, M., Chatterjee, D., Nayrac, M., et al. (2022). Strong humoral immune responses against SARS-CoV-2 Spike after BNT162b2 mRNA vaccination with a 16-week interval between doses. Cell Host Microbe 30, 97–109.e5. 10.1016/J.CHOM.2021.12.004/ATTACHMENT/BD4DA173-5A3F-408E-A5A2-9803CC3B4922/MMC1.PDF.

53. Kopr, M., Kobler, G., and Langhorne, J. (1994). Max·Planck·lnstitut fiir Immunbiologie, Freiburg P. chabaudi infection in ll..-4-deficient mice The immune response to Plasmodium chabaudi malaria in interleukin-4-deficient mice*.

54. Pape, K.A., Maul, R.W., Dileepan, T., Paustian, A.S., Gearhart, P.J., and Jenkins, M.K. (2018). Naïve B cells with high-avidity germline-encoded antigen receptors produce persistent IgM+ and transient IgG+ memory B cells. Immunity 48, 1135. 10.1016/J.IMMUNI.2018.04.019.

55. Chao, G., Lau, W.L., Hackel, B.J., Sazinsky, S.L., Lippow, S.M., and Wittrup, K.D. (2006). Isolating and engineering human antibodies using yeast surface display. Nat Protoc 1, 755–768. 10.1038/NPROT.2006.94.

56. Paus, D., Tri, G.P., Chan, T.D., Gardam, S., Basten, A., and Brink, R. (2006). Antigen recognition strength regulates the choice between extrafollicular plasma cell and germinal center B cell differentiation. J Exp Med 203, 1081. 10.1084/JEM.20060087.

57. Garrett Rappazzo, C., Tse, L. V., Kaku, C.I., Wrapp, D., Sakharkar, M., Huang, D., Deveau, L.M., Yockachonis, T.J., Herbert, A.S., Battles, M.B., et al. (2021). Broad and potent activity against SARS-like viruses by an engineered human monoclonal antibody. Science (1979) 371, 823–829. 10.1126/SCIENCE.ABF4830/SUPPL_FILE/ABF4830_TABLE_S2_GISAID_ACKNOWLEDGMENTS.PDF.

58. Turqueti-Neves, A., Otte, M., da Costa, O.P., Höpken, U.E., Lipp, M., Buch, T., and Voehringer, D. (2014). B-cell-intrinsic STAT6 signaling controls germinal center formation. Eur J Immunol 44, 2130–2138. 10.1002/eji.201344203.

59. Yusuf, I., Kageyama, R., Monticelli, L., Johnston, R.J., DiToro, D., Hansen, K., Barnett, B., and Crotty, S. (2010). Germinal Center T Follicular Helper Cell IL-4 Production Is Dependent on Signaling Lymphocytic Activation Molecule Receptor (CD150). The Journal of Immunology 185, 190–202. 10.4049/jimmunol.0903505.

60. King, I.L., and Mohrs, M. (2009). IL-4-producing CD4+ T cells in reactive lymph nodes during helminth infection are T follicular helper cells. Journal of Experimental Medicine 206, 1001–1007. 10.1084/jem.20090313.

61. Reinhardt, R.L., Liang, H.-E., and Locksley, R.M. (2009). Cytokine-secreting follicular T cells shape the antibody repertoire. Nat Immunol 10, 385–393. 10.1038/ni.1715.

62. Huse, M., Lillemeier, B.F., Kuhns, M.S., Chen, D.S., and Davis, M.M. (2006). T cells use two directionally distinct pathways for cytokine secretion. Nature Immunology 2006 7:3 7, 247–255. 10.1038/ni1304.

63. Koike, T., Harada, K., Horiuchi, S., and Kitamura, D. (2019). The quantity of CD40 signaling determines the differentiation of b cells into functionally distinct memory cell subsets. Elife 8. 10.7554/eLife.44245.001.

64. Chou, C., Verbaro, D.J., Tonc, E., Holmgren, M., Cella, M., Colonna, M., Bhattacharya, D., and Egawa, T. (2016). The Transcription Factor AP4 Mediates Resolution of Chronic Viral Infection through Amplification of Germinal Center B Cell Responses. Immunity 45, 570–582. 10.1016/j.immuni.2016.07.023.

65. Quast, I., Dvorscek, A.R., Pattaroni, C., Steiner, T.M., McKenzie, C.I., Pitt, C., O’Donnell, K., Ding, Z., Hill, D.L., Brink, R., et al. (2022). Interleukin-21, acting beyond the immunological synapse, independently controls T follicular helper and germinal center B cells. Immunity 0. 10.1016/J.IMMUNI.2022.06.020.

66. Iii, H.C.M., Lipsky, P.E., Leonard, W.J., Shaffer, D.J., Akilesh, S., Roopenian, D.C., Kim, H.-P., Wang, G., Qi, C.-F., Ozaki, P.K., et al. (2021). Inducer of Blimp-1 and Bcl-6 Plasma Cell Generation by IL-21, a Novel Regulation of B Cell Differentiation and. J Immunol 173, 5361–5371. 10.4049/jimmunol.173.9.5361.

67. Kaplan, M.H., Schindler, U., Smiley, S.T., and Grusby, M.J. (1996). Stat6 is required for mediating responses to IL-4 and for the development of Th2 cells. Immunity 4, 313–319. 10.1016/S1074-7613(00)80439-2.

68. Herbert, D.R., Hölscher, C., Mohrs, M., Arendse, B., Schwegmann, A., Radwanska, M., Leeto, M., Kirsch, R., Hall, P., Mossmann, H., et al. (2004). Alternative macrophage activation is essential for survival during schistosomiasis and downmodulates T helper 1 responses and immunopathology. Immunity 20, 623–635. 10.1016/S1074-7613(04)00107-4.

69. Barner, M., Mohrs, M., Brombacher, F., and Kopf, M. (1998). Differences between IL-4Rα-deficient and IL-4-deficient mice reveal a role for IL-13 in the regulation of Th2 responses. Current Biology 8, 669–672. 10.1016/s0960-9822(98)70256-8.

70. Mohrs, K., Wakil, A.E., Killeen, N., Locksley, R.M., and Mohrs, M. (2005). A two-step process for cytokine production revealed by IL-4 dual-reporter mice. Immunity 23, 419–429. 10.1016/j.immuni.2005.09.006.

71. Strasser, A., Whittingham, S., Vaux, D.L., Bath, M.L., Adams, J.M., Cory, S., and Harris, A.W. (1991). Enforced BCL2 expression in B-lymphoid cells prolongs antibody responses and elicits autoimmune disease. Proc Natl Acad Sci U S A 88, 8661. 10.1073/PNAS.88.19.8661.

72. Ndungu, F.M., Cadman, E.T., Coulcher, J., Nduati, E., Couper, E., MacDonald, D.W., Ng, D., and Langhorne, J. (2009). Functional Memory B Cells and Long-Lived Plasma Cells Are Generated after a Single Plasmodium chabaudi Infection in Mice. PLoS Pathog 5, e1000690. 10.1371/JOURNAL.PPAT.1000690.

73. Keitany, G.J., Kim, K.S., Krishnamurty, A.T., Hondowicz, B.D., Hahn, W.O., Dambrauskas, N., Sather, D.N., Vaughan, A.M., Kappe, S.H.I., and Pepper, M. (2016). Blood Stage Malaria Disrupts Humoral Immunity to the Pre-erythrocytic Stage Circumsporozoite Protein. Cell Rep 17, 3193–3205. 10.1016/j.celrep.2016.11.060.

74. Taylor, J.J., Martinez, R.J., Titcombe, P.J., Barsness, L.O., Thomas, S.R., Zhang, N., Katzman, S.D., Jenkins, M.K., and Mueller, D.L. (2012). Deletion and anergy of polyclonal B cells specific for ubiquitous membrane-bound self-antigen. Journal of Experimental Medicine 209, 2065–2077. 10.1084/JEM.20112272.

75. vander Heiden, J.A., Yaari, G., Uduman, M., Stern, J.N.H., O’connor, K.C., Hafler, D.A., Vigneault, F., and Kleinstein, S.H. (2014). pRESTO: a toolkit for processing high-throughput sequencing raw reads of lymphocyte receptor repertoires. Bioinformatics 30, 1930. 10.1093/BIOINFORMATICS/BTU138.

76. BBTools User Guide - DOE Joint Genome Institute https://jgi.doe.gov/data-and-tools/software-tools/bbtools/bb-tools-user-guide/.

77. Gupta, N.T., vander Heiden, J.A., Uduman, M., Gadala-Maria, D., Yaari, G., and Kleinstein, S.H. (2015). Change-O: a toolkit for analyzing large-scale B cell immunoglobulin repertoire sequencing data. Bioinformatics 31, 3356. 10.1093/BIOINFORMATICS/BTV359.

78. Wickham, H., Averick, M., Bryan, J., Chang, W., D’, L., Mcgowan, A., François, R., Grolemund, G., Hayes, A., Henry, L., et al. (2019). Welcome to the Tidyverse. J Open Source Softw 4, 1686. 10.21105/JOSS.01686.

