## Supplemental Information for "IL-4 downregulates BCL6 to promote memory B cell selection in germinal centers"

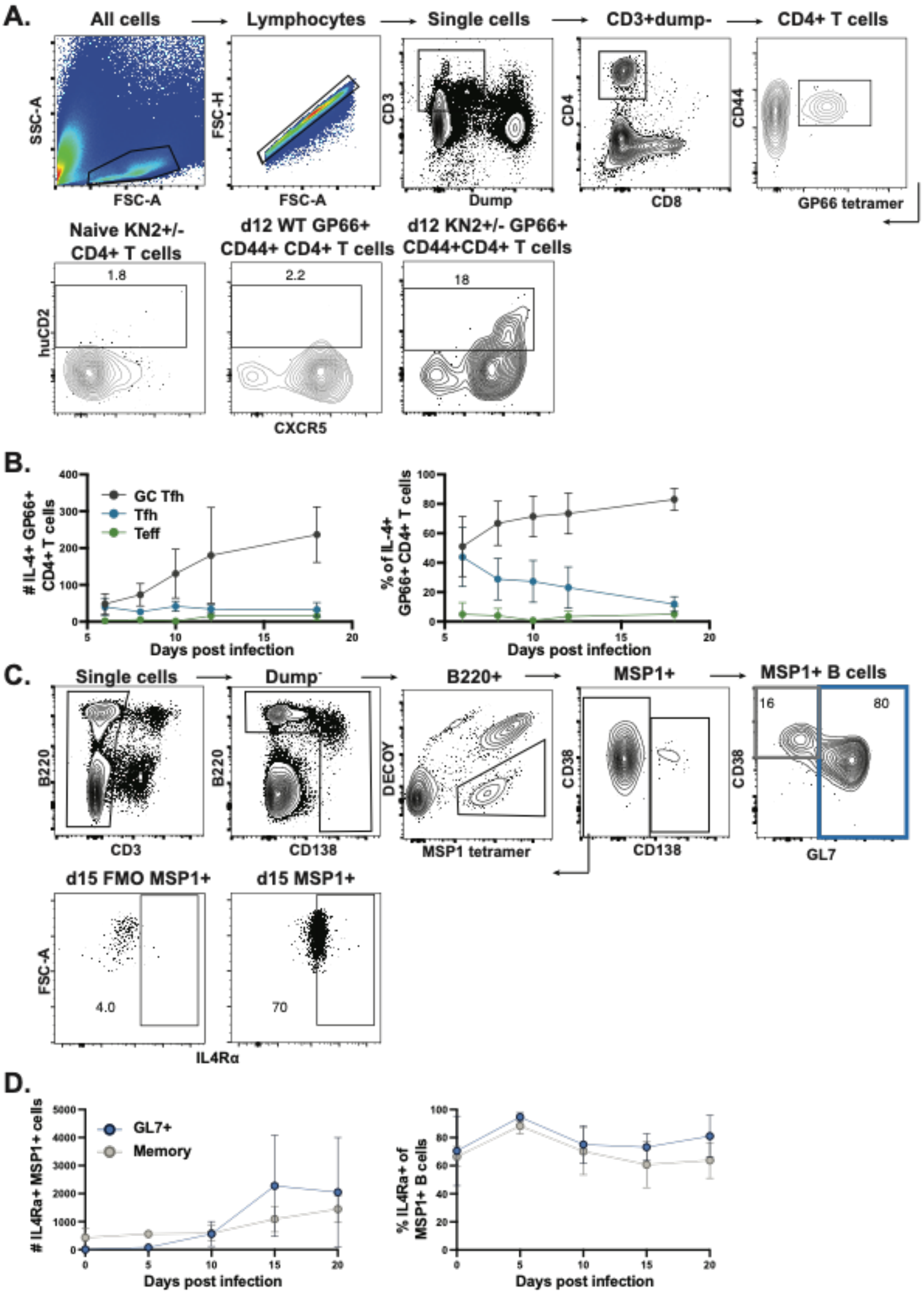

A.
All cells
Lymphocytes
Single cells
CD3+dump-
CD4+ T cells
SSC-A
FSC-A
FSC-H
FSC-A
CD3
Dump
CD4
CD8
CD44
GP66 tetramer
Naive KN2+/- CD4+ T cells
d12 WT GP66+ CD44+ CD4+ T cells
d12 KN2+/- GP66+ CD44+CD4+ T cells
1.8
2.2
18
huCD2
CXCR5
B.
GC Tfh
Tfh
Teff
### IL-4+ GP66+ CD4+ T cells
% of IL-4+ GP66+CD4+ T cells
Days post infection
Days post infection
C.
Single cells
Dump-
B220+
MSP1+
MSP1+ B cells
B220
CD3
B220
CD138
DECOY
MSP1 tetramer
CD38
CD138
16
80
CD38
GL7
d15 FMO MSP1+
d15 MSP1+
FSC-A
4.0
70
IL4Rα
D.
GL7+
Memory
### IL4Ra+ MSP1+ cells
% IL4Ra+ of MSP1+ B cells
Days post infection
Days post infection

**Figure S1. Kinetics of IL-4 production and IL4Rα expression throughout acute *Plasmodium* infection. Related to Figure 1.** $KN2^{+/-}$ mice were infected with blood-stage *P.y*-GP66, and GP66-specific $CD4^+$ T cells and MSP1-specific B cells were assessed for IL-4 production or IL4Ra expression, respectively, at the time points indicated. (**A**) Representative gating strategy for evaluating GP66-specific $CD4^+$ T cells. Non-doublet lymphocytes were gated on $CD3^+B220^-CD11b^-CD11c^-CD4^+CD8^-$, then further gated as $CD44^+GP66^+$. Gating on huCD2 expression on naive $KN2^{+/-}$ $CD4^+$ T cells (left), and WT (middle) and $KN2^{+/-}$ (right) $gp66^+CD44^+CD4^+$ T cells 12 days post-*P.y*-GP66 infection. (**B**) Number (left) and frequency (right) of IL-4-producing ($huCD2^+$) GP66-specific $CD4^+$ T cells by subset, with GC Tfh identified as $PD\text{-}1^+CXCR5^-$ (grey), Tfh as $PD\text{-}1^-CXCR5^+$ (blue), and Teff as $PD\text{-}1^-CXCR5^-$ (green). Data are combined from three independent experiments with 6-8 mice per time point. (**C**) Representative gating strategy for evaluating MSP1-specific B cells and plasmablasts. Non-doublet lymphocytes (A) were further gated on $CD3^-B220^+decoy^-MSP1^+$ cells. Memory B cells (grey) were defined as $CD138^-CD38^+GL7^-$ and $GL7^+$ B cells (blue) as $CD138^-CD38^{+/-}GL7^+$. Representative gating for IL4Rα on MSP1-specific B cells from WT mice infected with *P.y*-GP66 using a staining panel excluding (left; fluorescence minus one or FMO) or including (right) IL4Rα. (**D**) Number and frequency of IL4Rα-expressing $MSP1^+$ memory and $GL7^+$ B cells. Data are combined from three independent experiments with 6 mice per time point.

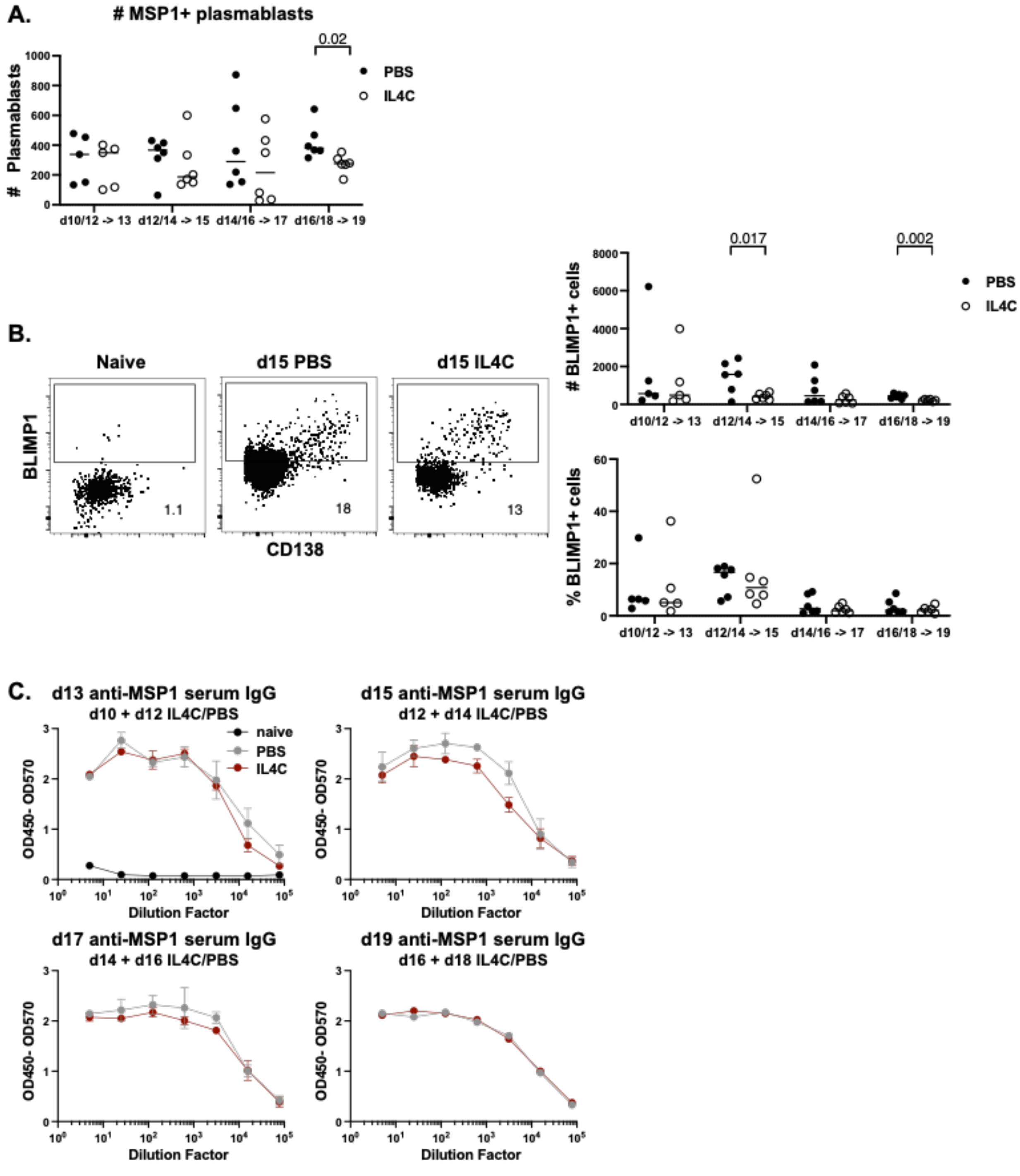


**Figure S2. IL4C treatment does not impact plasma cell differentiation. Related to Figure 1.** WT mice were infected with *P.ch* and treated with PBS or IL4C for two-day treatment windows as shown in Fig 1D. (**A**) Number of MSP1$^+$ plasmablasts (gating as in fig. S1C) at each endpoint. (**B**) Representative flow plots and quantification of BLIMP1 expression in total MSP1$^+$B220$^+$CD3$^-$ lymphocytes in naive mice, or mice treated with PBS or IL4C on days 12 and 14 and analyzed on day 15. (**C**) Serum was collected at each endpoint and anti-MSP1 IgG levels were measured by ELISA. Anti-MSP1 IgG ELISA dilution curves from serum collected at each endpoint. Data are combined from two independent experiments with 5-6 mice per group.

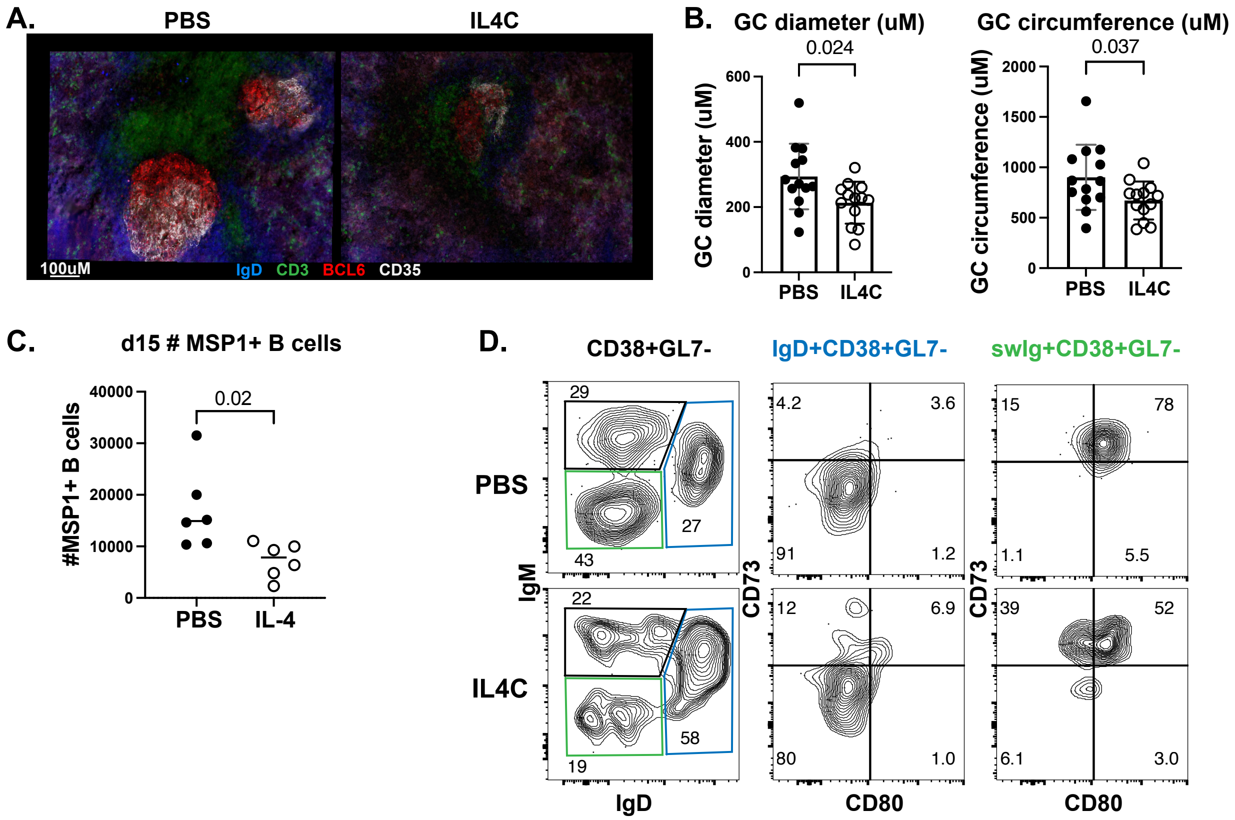


**Figure S3. IL-4 administered after GC formation limits GC size and output. Related to Figure 2.** WT mice were infected with *P.ch* and treated with IL4C or PBS on days 12 and 14, and analyzed on day 15. **(A)** Representative images of GCs from spleen sections stained with anti-IgD, anti-CD3, anti-BCL6, and anti-CD35; PBS-treated (left) and IL4C-treated (right). (**B**) Diameter (left) and circumference (right) of GCs in mice treated with PBS or IL4C; each dot represents one GC. Spleens were collected from two independent experiments and 3 spleens per group were sectioned and stained, with 4-5 GCs analyzed per section. (**C**) Number of MSP1$^{+}$ B cells at day 15 in mice treated with PBS or IL-4 on days 12, 13, and 14 post-*P.ch* infection. Data are representative of two experiments with 6 mice per group. (**D**) Representative flow cytometry plots indicating gating of CD73 and CD80 on MSP1-specific IgD$^{+}$ (blue) and swIg$^{+}$ (green) CD38$^{+}$GL7$^{-}$ B cells on day 15 in PBS- or IL4C-treated mice.

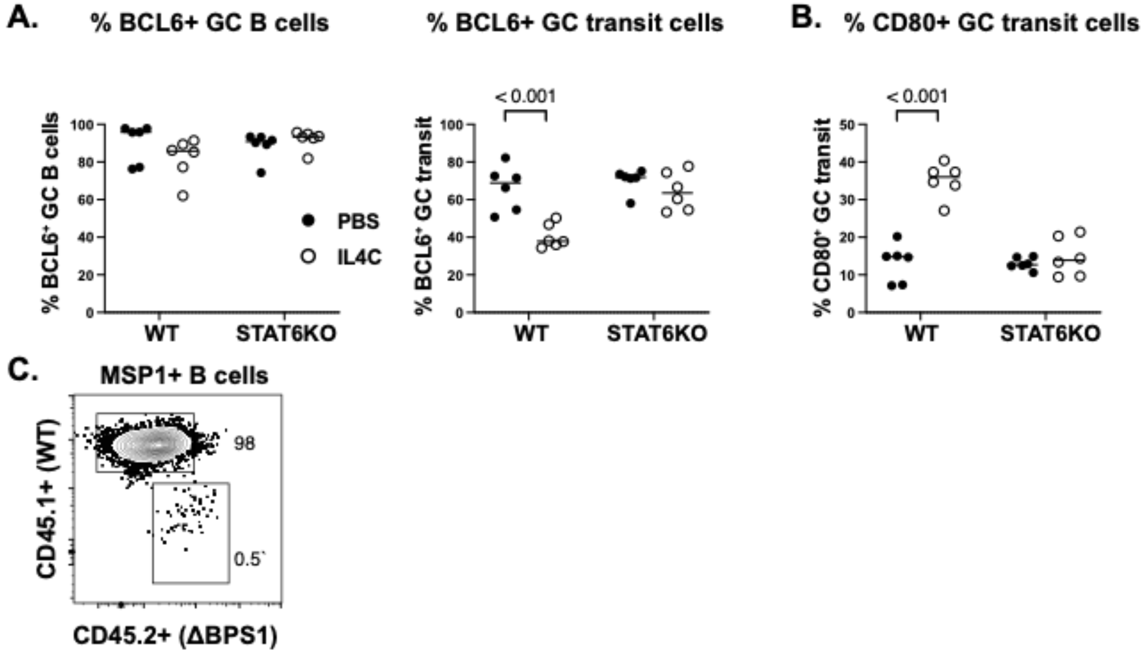


**Figure S4. IL-4-mediated acts directly on B cells in a STAT6-dependent manner to downregulate BCL6. Related to Figure 3.** WT and STAT6KO mice were infected with *P.ch* and treated with IL4C or PBS on days 12 and 14, and the MSP1-specific B cell response was analyzed on day 15. (**A**) Frequency of MSP1$^{+}$ GC B cells and GC transit B cells expressing BCL6. (**B**) Frequency of MSP1$^{+}$ GC transit B cells expressing CD80. Data are combined from two independent experiments with 6 mice per group. (**C**) Splenic B cells were isolated from ΔBPS1 mice and transferred into WT hosts. The mice were infected with *P.ch*, treated with IL4C on days 12 and 14, and the MSP1-specific B cell response was analyzed on day 15. Representative flow plot indicating gating on MSP1$^{+}$CD45.1$^{+}$ WT and CD45.2$^{+}$ ΔBPS1 B cells.

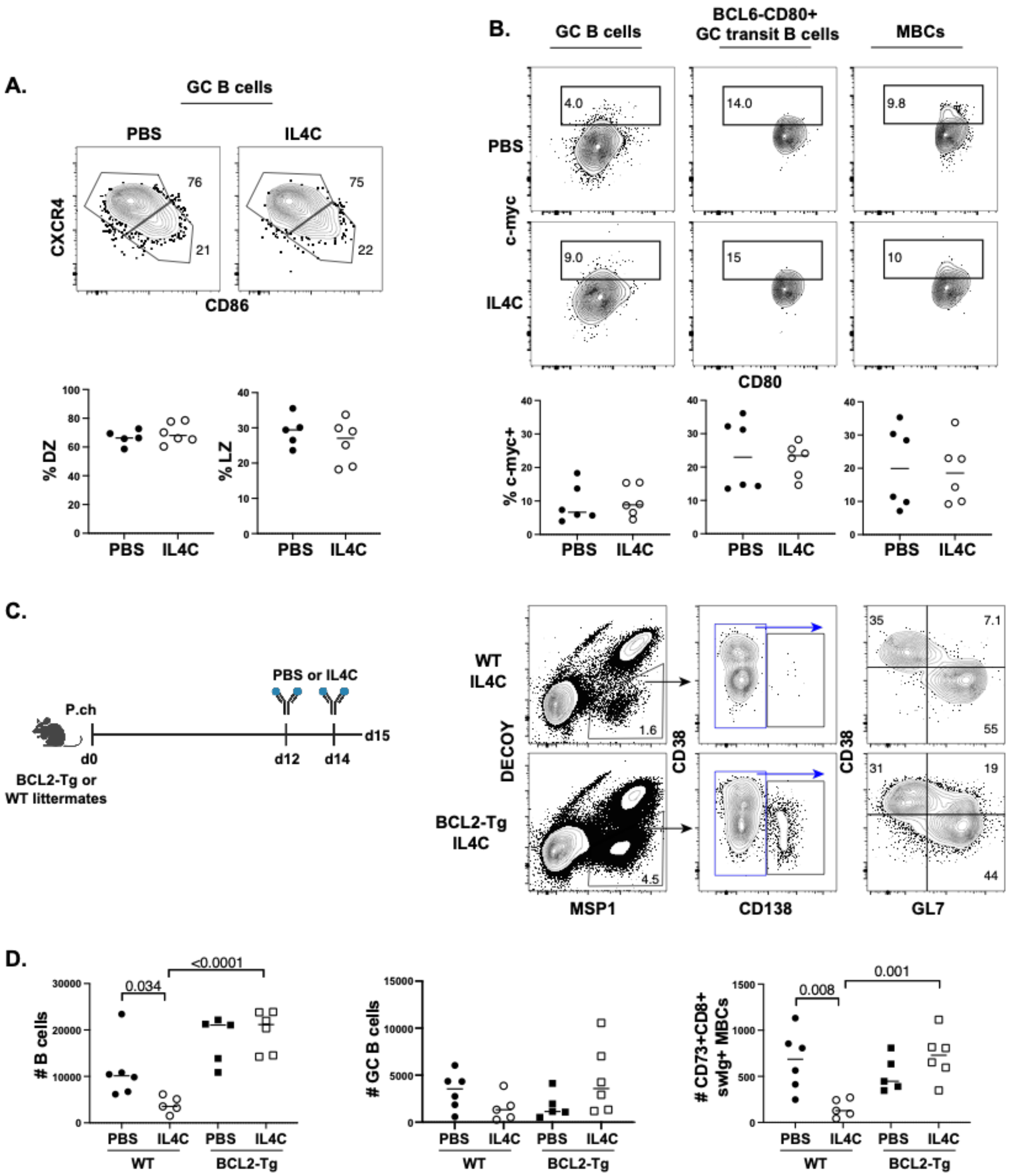


**Figure S5. IL-4 increases pre-memory GC B cell death. Related to Figure 4.** WT mice were infected with *P.ch* and treated with PBS or IL4C on days 12 and 14. (**A**) Representative flow plots and quantification of MSP1$^+$ GC B cells in the dark zone (CXCR4$^{hi}$CD86$^{lo}$) or light zone (CXCR4$^{lo}$CD86$^{hi}$). (**B**) Representative flow plots and quantification of MSP1$^+$c-myc$^+$ GC B cells (left), BCL6$^-$CD80$^+$ GC transit B cells (middle), and MBCs (right). Data are combined from three independent experiments with 5-6 mice per group. (**C**) WT mice and BCL2-Tg littermates were infected with *P.ch* and treated with PBS or IL4C on days 12 and 14 and analyzed on day 15.

Representative flow plots showing gating on $MSP1^+$ B cells from WT (top) and BCL2-Tg (bottom) mice treated with IL4C. (**D**) Number of $MSP1^+$ total B cells, GC B cells, and $swIg^+$ $CD73^+CD80^+$ MBCs in each group. Data are combined from three independent experiments with 5-6 mice per group.

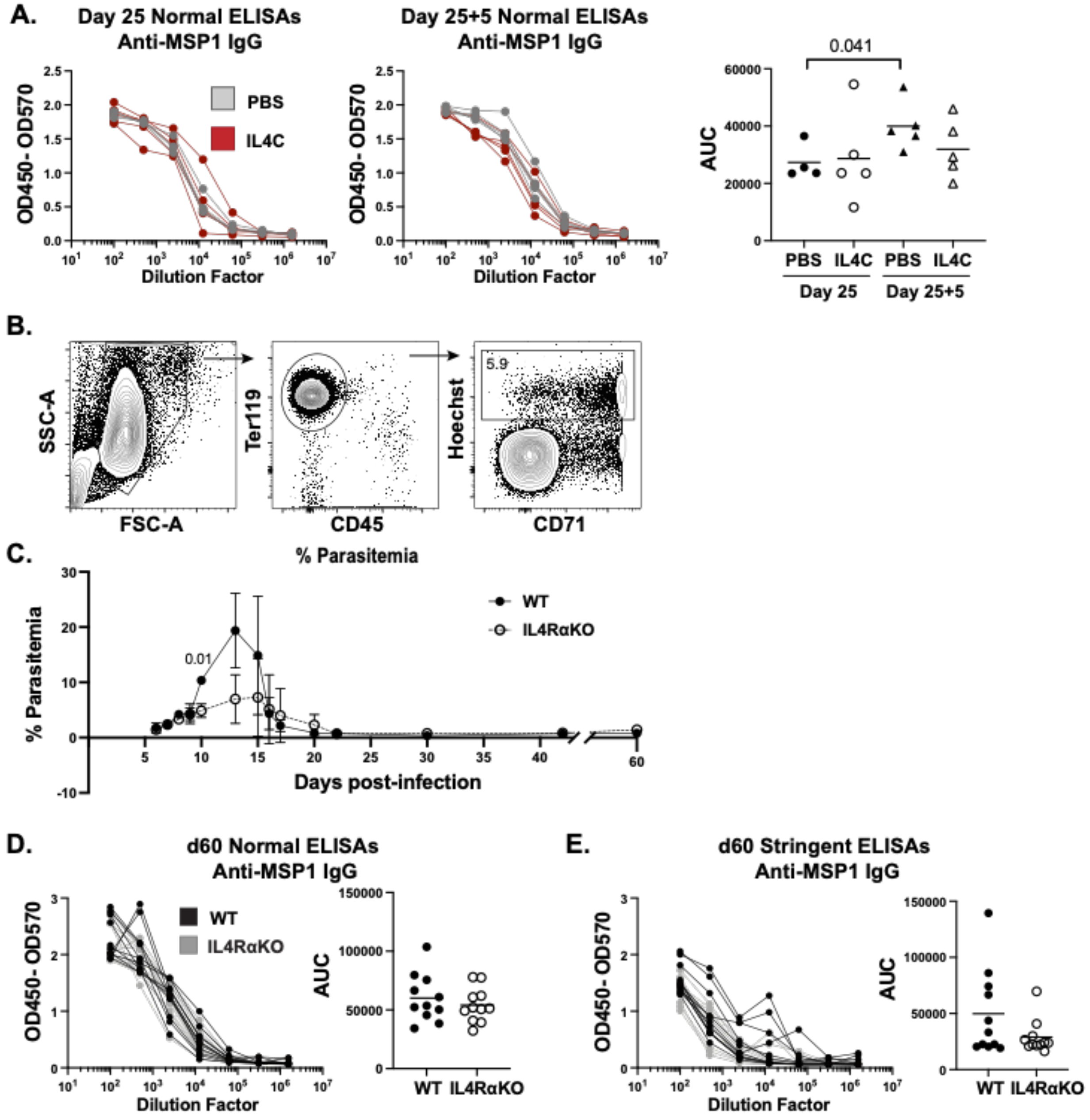


**Figure S6. Characterization of the memory response in IL4RαKO mice. Related to Figure 5.** WT mice were infected with *P.y.*-GP66 and treated with PBS or IL4C every other day between days 12-25, and re-challenged on day 25. (**A**) Normal ELISA dilution curves and AUC for anti-MSP1 IgG in serum collected at day 25 and 5 days post-challenge. Data are combined from two independent experiments with 4-5 mice per group. WT and IL4RαKO mice were infected with *P.y*-GP66 and analyzed 60 days post-infection. (**B**) Representative gating scheme to identify *Plasmodium*-infected iRBCs, gated as Ter119$^+$CD45$^-$Hoechst$^+$ in 1uL of blood from a WT mouse 9 days post-infection. (**C**) Parasitemia throughout the 60 days of infection in each group. Data are combined from three independent experiments with 6-7 mice per group. Dilution curves and AUC of (**E**) normal and (**F**) stringent ELISAs to assess anti-MSP1 IgG serum levels. Data are combined from four independent experiments with 11 mice per group.

| Surface molecule | Fluorochrome | Clone | Manufacturer |
|---|---|---|---|
| B220 | BV510 | RA3-6B2 | BD Biosciences |
| B220 | BUV737 | RA3-6B2 | BD Biosciences |
| B220 | PE-CF594 | RA3-6B2 | BD Biosciences |
| BCL2 | AF488 | BCL/10C4 | BioLegend |
| BCL6 | AF647 | K112-91 | BD Biosciences |
| BCL6 | PE-CF594 | K112-91 | BD Biosciences |
| BLIMP1 | PE-CF594 | 6D3 | BD Biosciences |
| CD3 | APC-ef780 | 145-2C11 | Invitrogen |
| CD3 | PerCP-Cy5.5 | 145-2C11 | BD Biosciences |
| CD4 | BUV805 | GK1.5 | BD Biosciences |
| CD4 | BV711 | RM4-5 | BioLegend |
| CD8 | BV510 | 53-6.7 | BD Biosciences |
| CD11b | PE-CF594 | M1/70 | BD Biosciences |
| CD11c | PE-CF594 | HL3 | BD Biosciences |
| CD23 | BV711 | B3B4 | BD Biosciences |
| CD38 | AF700 | 90 | Invitrogen |
| CD44 | AF700 | IM7 | BD Biosciences |
| CD45.1 | FITC | A20 | eBioscience |
| CD45.2 | APC-ef780 | 104 | Invitrogen |
| CD62L | BV786 | MEL-14 | BD Biosciences |
| CD73 | PE-Cy7 | TY/11.8 | Invitrogen |
| CD80 | BV605 | 16-10A1 | BD Biosciences |
| CD86 | BV605 | GL-1 | BioLegend |
| CD138 | BB515 | 281-2 | BD Biosciences |
| CD138 | BV650 | 281-2 | BD Biosciences |
| c-myc | FITC | Y69 | abcam |
| CXCR4 | BV711 | L276F12 | BioLegend |
| CXCR5 | PE | 2G8 | BD Biosciences |
| GL7 | ef450 | GL-7 | Invitrogen |
| huCD2 | Biotin | RPA-2.10 | Invitrogen |
| IgM | BV786 | II/41 | BD Biosciences |
| IgD | BUV395 | 11-26c.2a | BD Biosciences |
| IL4R$\alpha$ | Biotin | mIL4R-M1 | BD Biosciences |
| IRF4 | FITC | 3E4 | Invitrogen |
| PD-1 | ef450 | J43 | Invitrogen |
| PD-1 | PE-Cy7 | 29F.1A12 | BioLegend |
| Streptavidin | BV605 | --- | BD Biosciences |
| Streptavidin | BV711 | --- | BD Biosciences |
| Streptavidin | BUV661 | --- | BD Biosciences |

**Table S1. Antibodies used for flow cytometry. Related to STAR Methods.**

| Adapter | Sequence |
|---|---|
| 1 Top | 5'-GTGACTGGAGTTCAGACGTGTGCTCTTCCGATCTNNNNNNNNNNNNNNNNNACACTACTCG*T-3' |
| 1 Bottom | 5'-/5Phos/CGAGTAGTGT-3' |
| 2 Top | 5'-GTGACTGGAGTTCAGACGTGTGCTCTTCCGATCTNNNNNNNNNNNNNNNNNTGTGCGGCTC*T-3' |
| 2 Bottom | 5'-/5Phos/GAGCCGCACA-3' |

**Table S2. UMI adapters for BCR sequencing. Related to STAR Methods.** Duplexed by IDT.

**Data S1. Related to Figure 5.** Sequencing data for BCRs cloned and sequenced from $CD80^+$ GC transit B cells and $CD73^+CD80^+$ MBCs sorted from mice treated with PBS or IL4C.
